# Dorsal raphe neurons signal integrated value during multi-attribute decision-making

**DOI:** 10.1101/2023.08.17.553745

**Authors:** Yang-Yang Feng, Ethan S. Bromberg-Martin, Ilya E. Monosov

## Abstract

The dorsal raphe nucleus (DRN) is implicated in psychiatric disorders that feature impaired sensitivity to reward amount, impulsivity when facing reward delays, and risk-seeking when grappling with reward uncertainty. However, whether and how DRN neurons signal reward amount, reward delay, and reward uncertainty during multi-attribute value-based decision-making, where subjects consider all these attributes to make a choice, is unclear. We recorded DRN neurons as monkeys chose between offers whose attributes, namely expected reward amount, reward delay, and reward uncertainty, varied independently. Many DRN neurons signaled offer attributes. Remarkably, these neurons commonly integrated offer attributes in a manner that reflected monkeys’ overall preferences for amount, delay, and uncertainty. After decision-making, in response to post-decision feedback, these same neurons signaled signed reward prediction errors, suggesting a broader role in tracking value across task epochs and behavioral contexts. Our data illustrate how DRN participates in integrated value computations, guiding theories of DRN in decision-making and psychiatric disease.

## Introduction

The dorsal raphe nucleus (DRN), the main source of serotonin to the forebrain^1^, is strongly implicated in various psychiatric disorders associated with deficits in value-based decision-making^2–11^. Many of these deficits, including impaired sensitivity to reward amount^12,13^, impulsivity when facing reward delays^14–23^, and risk-seeking when grappling with reward uncertainty^12,24–26^ can be experimentally recapitulated by manipulating serotonergic transmission or DRN activity. However, while DRN neurons have been shown to be sensitive to a diverse range of behavioral paradigms and task variables in rodents^10,11,27–32^, much less is known about DRN’s functional roles in primates^33–36^. Further, little is known about DRN’s role in complex cognitive tasks like multi-attribute value-based decision-making, where subjects must integrate reward, delay, and uncertainty to judge an offer’s overall subjective value and make a choice.

Despite this gap in knowledge, there are several theories about the functional role of DRN that have key implications for clinical intervention. One important set of theories suggests that DRN neurons broadcast ‘beneficialness’^11,37^ or ‘the availability of time and resources’^10^, a signal that integrates multiple distinct features or attributes to reflect the overall subjective value of offers or motivational outcomes and guide the neural computations that underlie emotion, mood, and value-guided behaviors, including multi-attribute decision-making. If so, DRN neurons should signal the subjective value of multi-attribute offers, for example, by integrating over reward amount, delay, and uncertainty. Another important set of theories suggests that DRN is involved in continuously tracking value-related statistics on multiple timescales, in the service of motivating behavior^6,27,29,31,33,35,38^. However, much is still unknown about whether and how DRN neurons signal such motivational variables, including surprise^28,39^, signed prediction errors^2^, or ‘global reward state’ (long-term average reward)^2,18,29,40,41^, particularly in the context of primate decision-making.

To test how DRN neurons process reward, delay, and uncertainty during decision-making, we recorded single DRN neurons from two monkeys as they chose between offers whose attributes, namely expected reward amount, reward delay, and reward uncertainty, varied independently. We found that many DRN neurons signaled offer attributes. Remarkably, these neurons commonly integrated offer attributes in a manner that reflected monkeys’ overall preferences for amount, delay, and uncertainty. After decision-making, in response to post-decision feedback, these same neurons signaled signed reward prediction errors, suggesting a broader role in tracking value across task epochs and behavioral contexts. Our data illustrate how DRN participates in integrated value computations, guiding theories of DRN in decision-making and psychiatric disease.

## Results

### Monkeys integrate reward amount, delay, and uncertainty to make multi-attribute decisions

We recorded single DRN neurons from two monkeys as they chose between two multi-attribute offers that each varied independently in expected reward amount (E[R]), reward delay (T[R]), and reward uncertainty (Unc[R]) (Figure 1A). Each offer, if chosen, provided a juice reward that was randomly drawn from a set of four possible outcomes, depicted as four bars of juice (Figure 1B, left). Thus, different offers provided different probability distributions of rewards, which could have different levels of E[R] (mean height of the bars) and Unc[R] (variability of the height of the bars, operationalized here as standard deviation). T[R], the delay between the time of choice and time of reward delivery, also varied across offers and was indicated by a ‘visual clock’ that provided explicit, online timing of progress towards reward delivery (Figure 1B, right).

**Figure 1.**
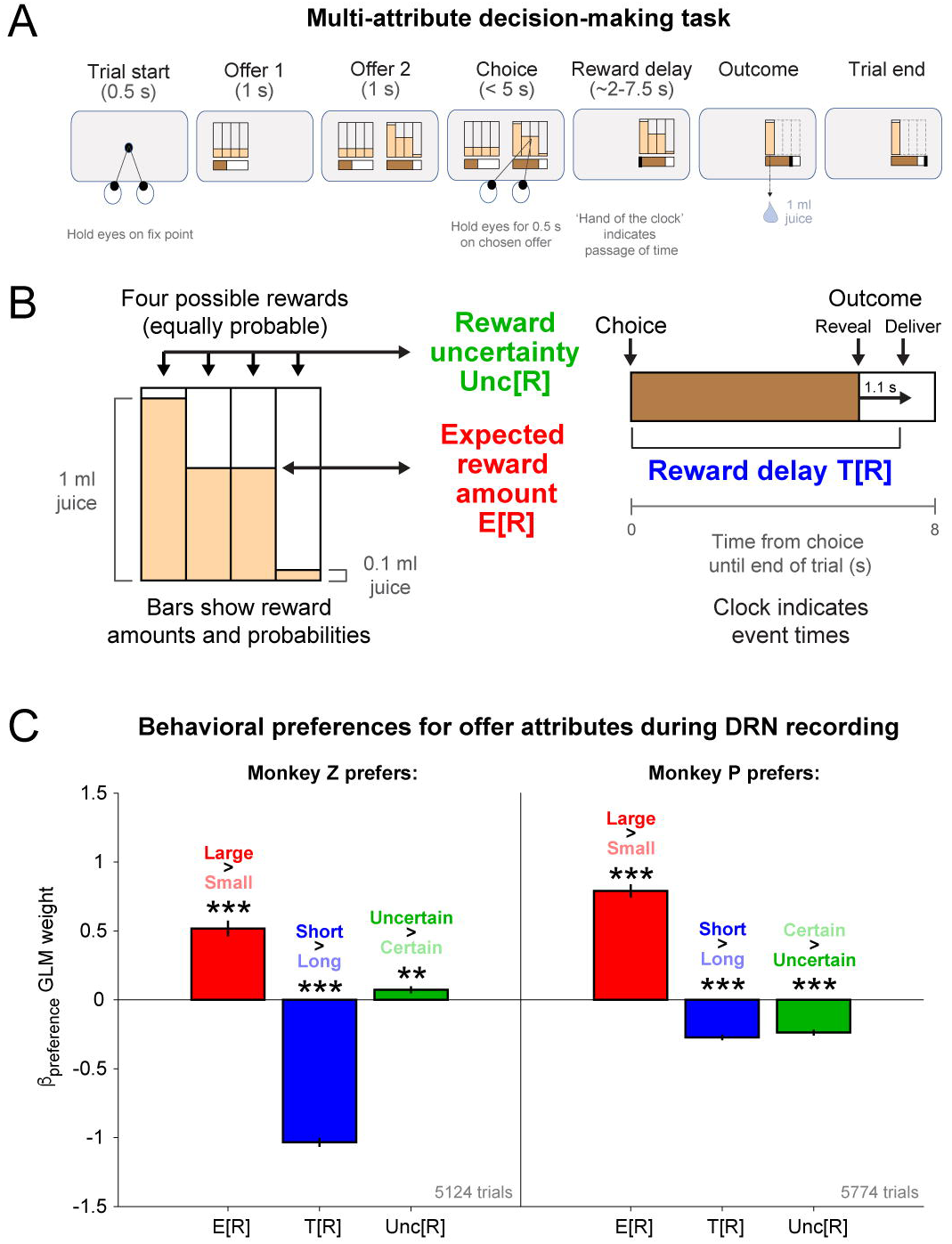
Monkeys exhibit preferences for expected reward, delay, and uncertainty during multi- attribute decision-making. (A) Two offers (Offer 1 and Offer 2) are presented sequentially and then the monkey has 5 seconds to free view and make a choice by fixating one offer for 0.5 seconds. After a delay, the outcome is revealed and the reward is delivered. (B) Left: Vertical bars represent the probability distribution over rewards, indicating the expected reward amount (E[R]) and reward uncertainty (Unc[R]). Right: Horizontal bar acts as a clock, representing reward delay (T[R]). To allow monkeys to prepare to consume the upcoming reward, when the reward delay has elapsed, the stimulus changes to reveal the outcome, and then the reward is delivered 1.1 s later (Methods). The total time between choice and end of the trial was always 8s. (C) Fitted weights describing each monkey’s subjective preferences for E[R], T[R], and Unc[R]. Error bars denote s.e.m. Both monkeys placed significant weights on each reward attribute (** p<0.01, *** p<0.001).

To quantify the effect of reward attributes on monkeys’ decision-making, we employed a standard approach of fitting their choice behavior using a generalized linear model (GLM) where the log-odds of choosing between the two offers is equal to the difference in their overall subjective values, and where the overall subjective value of each offer is equal to a linear combination of offer attributes multiplied by their respective weights on value (Methods). Examining the resulting weights revealed that both monkeys understood the task and its offers. Consistent with previous work^42–44^, monkeys chose offers more when they had larger E[R] and shorter T[R] (Figure 1C). The monkeys also had significant preferences for Unc[R], though in different manners, again consistent with individual variability in risk preferences seen in our^42,45^ and others’^46,47^ previous work.

Thus, monkeys demonstrated preferences for expected reward amount, reward delay, and reward uncertainty as they traded off these attributes to make multi-attribute decisions. We next asked whether and how DRN neurons signal these attributes during multi-attribute decision-making.

### DRN neurons signal reward amount, delay, and uncertainty during multi-attribute decision-making

We recorded 219 DRN neurons as the two monkeys performed the task (n=117 in Monkey Z, and n=102, in Monkey P). The recording locations were verified using MRI imaging of the recording electrode within DRN (Figure 2A) and histology (Figure 2B). We reconstructed the neurons’ positions within DRN using previously established methods (Figure 2C). We also verified that our recordings took place in a subregion of DRN that is dense in serotonergic cell bodies^48–50^, by examining our MRI- and histology-based reconstruction along with macaque DRN tissue that was stained for serotonin transporter (SERT) using previously established methods^51^ (Figure 2D).

**Figure 2.**
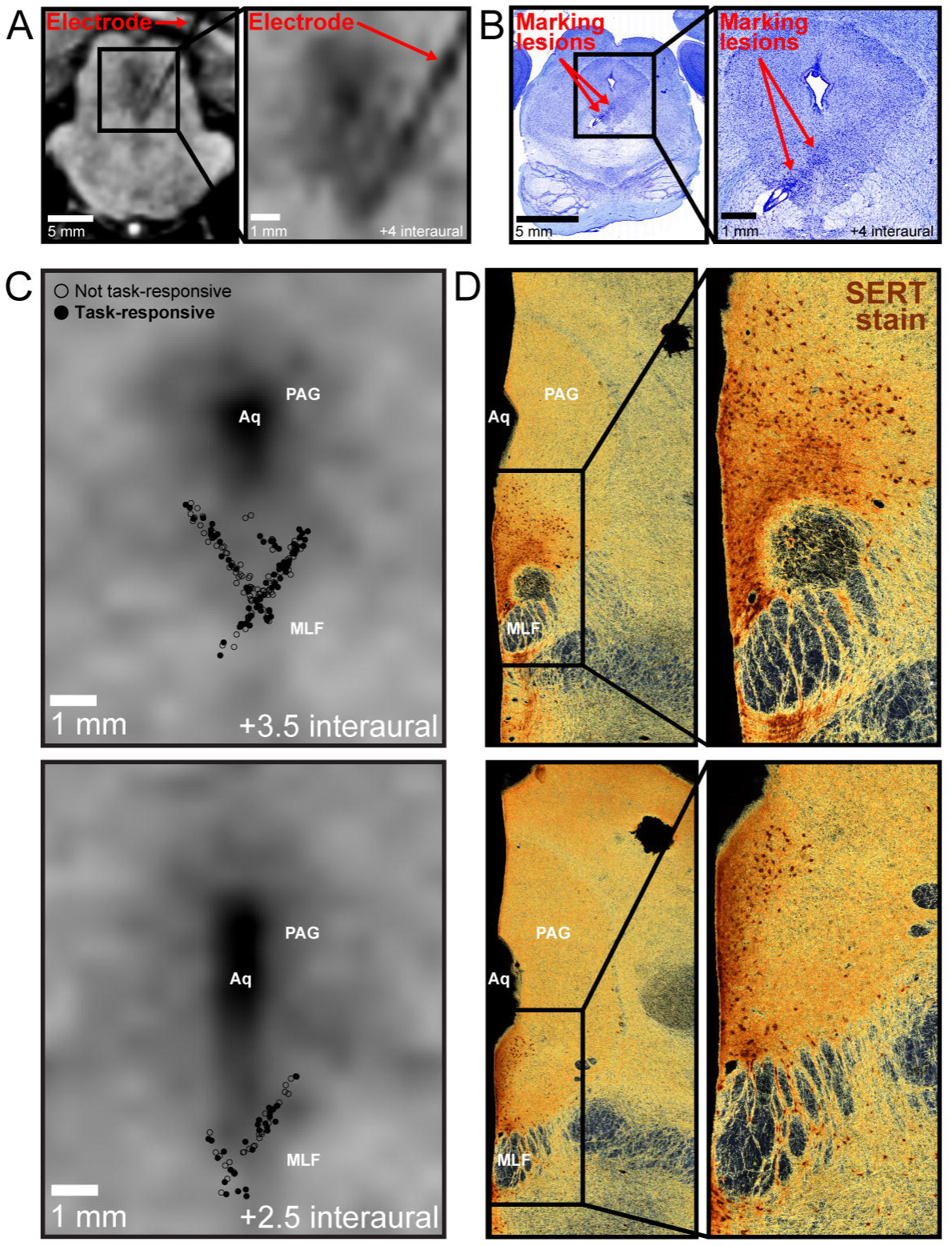
Recording single neurons in primate dorsal raphe nucleus. (A) MRI taken with electrode in recording location, verifying its position in DRN. Shown are coronal views slightly tilted to align with the electrode track. The electrode is visible as a black ‘shadow’ on the MRI. (B) Nissl staining showing electrode track and marking lesions (Red arrows), delineating approximate range of recordings. The trochlear nucleus is visible just ventral to the recording track. (C) Anatomical reconstruction of recorded neurons, based on previously published methods^238^. Empty circles represent recorded neurons that were not task-responsive. Filled circles are task-responsive neurons. Top: Neurons from +3 to +4.5 interaural are shown on a plane corresponding to approximately +3.5 interaural. Bottom: Neurons from +2 to +2.5 interaural are shown on a plane corresponding to approximately +2.5 interaural. (D) SERT staining of tissue slices cut along the transverse axis of the brainstem, approximately corresponding to reconstructions shown in (C). Aq, cerebral aqueduct; MLF, medial longitudinal fasciculus; PAG, periaqueductal gray.

An analysis of variance across all epochs in our task, including trial start, choice, and post-decision epochs (Methods) revealed that 107/219 DRN neurons displayed task-related modulation. Neurons’ intrinsic electrophysiological properties and anatomical positions were not related to whether they were task-responsive (Figure 2C, Supplementary Figure 1, Supplementary Tables 1,2). We therefore focused our analyses on the 107 task-responsive DRN neurons.

We inspected the offer responses of DRN neurons and found that neurons commonly scaled their activity based on E[R], T[R], and Unc[R]. An example neuron’s responses to these offer attributes are shown in Figure 3A. Following offer presentation, this neuron displayed relatively phasic activations that scaled with E[R] (left), T[R] (middle), and Unc[R] (right). The mean cross-validated normalized activity (Methods) across task-responsive neurons demonstrated similar phasic dynamics that robustly discriminated offers based on their E[R], T[R], and Unc[R] (Figure 3B). To identify attribute-signaling neurons, we employed a standard approach of fitting neurons’ offer responses using a GLM where normalized firing rate is modeled as a linear weighted combination of offer attributes (Methods). Thus, a significant fitted weight for an attribute indicates that a neuron’s signals were related to that attribute. This approach identified significant proportions of task-responsive DRN neurons that signaled E[R], T[R], and Unc[R] (E[R]: 24/107, p=5.1E-10; T[R]: 22/107, p=1.5E-8; Unc[R]: 11/107, p=0.018; one-tailed binomial tests) (Figure 3C). Thus, all three of these offer attributes were signaled by DRN neurons.

**Figure 3.**
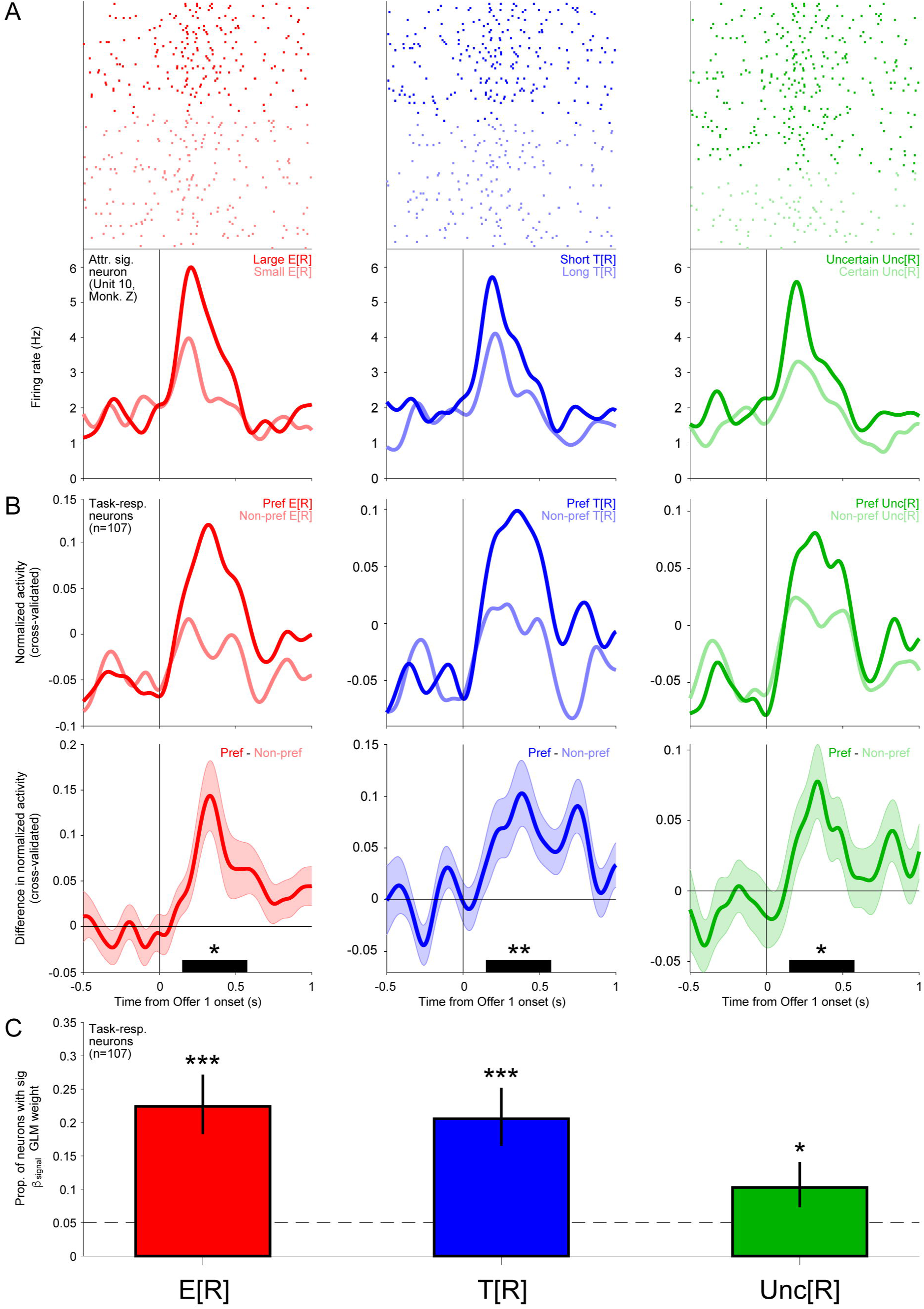
DRN neurons signal expected reward, delay, and uncertainty during multi-attribute decision- making. (A) Rasters and SDFs showing example neurons’ responses to Offer 1 on trials where it had Large vs Small E[R] (left, red), Short vs Long T[R] (center, blue), or Uncertain vs Certain Unc[R] (right, green). (B) Mean cross-validated normalized activity for task-responsive DRN neurons on trials where Offer 1 E[R] (left, red), T[R] (center, blue), or Unc[R] (right, green) was preferred or non-preferred by the neuron. (C) Difference in the activity shown in B. Error bars denote s.e.m. Average discrimination during the offer response analysis window (150-575 ms after Offer 1 presentation, black bar) was significant for E[R], T[R], and Unc[R] signals (two-tailed signed-rank test, *** p<0.001). (D) Proportion of task-responsive DRN neurons with significant fitted weights for each attribute, fitting normalized activity in response to Offer 1 (one-tailed binomial test, ** p<0.01, *** p<0.001). Error bars indicate 68% confidence interval (Clopper–Pearson method).

### DRN neurons integrate reward amount, delay, and uncertainty during multi-attribute decision-making

If DRN neurons signal the overall value of each offer, then they should signal attributes in accordance with each monkey’s subjective preferences, integrating these attributes by signaling their values jointly in the same direction as these preferences.

We therefore asked whether and how DRN neurons’ E[R], T[R], and Unc[R] signals reflected the subjective value monkeys assigned to each attribute based on their preferences. To test this, we quantified each task-responsive neuron’s signal for each attribute using an attribute value index, with indices above 0 indicating higher firing in response to offers with higher value based on that attribute (Methods). Strikingly, we found that all three attributes were predominantly signaled in the direction of subjective value (Figure 4). E[R]-signaling, T[R]-signaling, and Unc[R]-signaling neurons had significantly positive E[R], T[R], and Unc[R] value indices, respectively (E[R]: 0.23, p=8.1E-5; T[R]: 0.14, p=0.00098; Unc[R]: 0.16, p=0.0098; two-tailed signed-rank tests). Even considering all task-responsive neurons, the mean E[R] and T[R] value indices were significantly positive, and Unc[R] had a similar trend (E[R]: 0.068, p=6.4E-7; T[R]: 0.042, p=0.0047; Unc[R]: 0.019, p=0.10; two-tailed signed-rank tests). Moreover, for each attribute, there were significant proportions of neurons with significantly positive attribute value indices (E[R]: n=22/107, p=2.8E-14; T[R]: n=17/107, p=1.6E-9; Unc[R]: n=9/107, p=0.0015; two-tailed binomial tests), but not with significantly negative indices (E[R]: n=2/107, p=1; T[R]: n=5/107, p=0.20; Unc[R]: n=2/107, p=1; two-tailed binomial tests). Thus, task-responsive DRN neurons tended to signal attributes in accordance with monkeys’ preferences.

**Figure 4.**
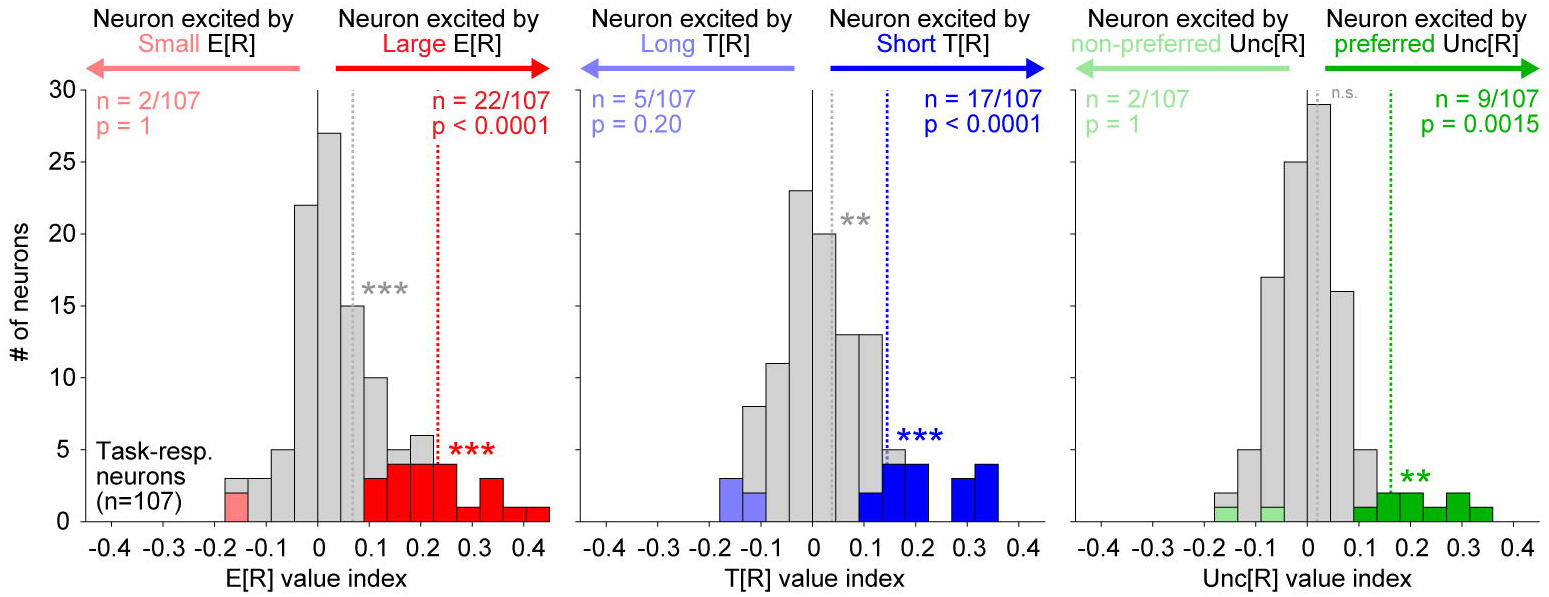
DRN neurons positively signal the value of expected reward, delay, and uncertainty. Histograms showing E[R] (left, red), T[R] (center, blue), Unc[R] (right, green) value indices (see Methods) of task-responsive neurons. Signals are biased in the direction of monkey preferences (dashed lines show mean; median significantly different from 0 by signed-rank test). The proportions of neurons with significant signaling of each attribute in the direction of monkey preferences are significant, while the proportions signaling the opposite are not (two-tailed binomial test). One outlier with an extreme E[R] value index was omitted from the left plot for presentation purposes (E[R] value index = 1.0, sig.). Colors indicate that neither (black), the x- coordinate (red), the y-coordinate (blue), or both (magenta) are significant (p < 0.05).

Next, we asked whether these attribute value signals are integrated. Specifically, we wondered whether DRN neurons’ E[R] signals, which convey information about an offer’s value based on its E[R], also conveyed information about an offer’s value based on its T[R] and Unc[R]. Consistent with integrated signaling of attributes, E[R] value indices were strongly correlated with T[R] and Unc[R] value indices, across task-responsive neurons (Figure 5A). Moreover, E[R]-signaling neurons strongly signaled T[R] and Unc[R] in the same direction as E[R] and with phasic dynamics that resembled how they signaled E[R] (Figure 5B). We also observed more neurons than expected by chance that had significant E[R] and T[R] value indices, or E[R] and Unc[R] value indices, in the same direction (E[R] and T[R]: n=10/107, p=3.3E-16; E[R] and Unc[R]: n=6/107, p=6.2E-09; two-tailed binomial tests, null proportion = 0.05*0.05/2). Thus, DRN neurons tended to integrate attributes, signaling them jointly and in directions consistent with monkeys’ preferences.

**Figure 5.**
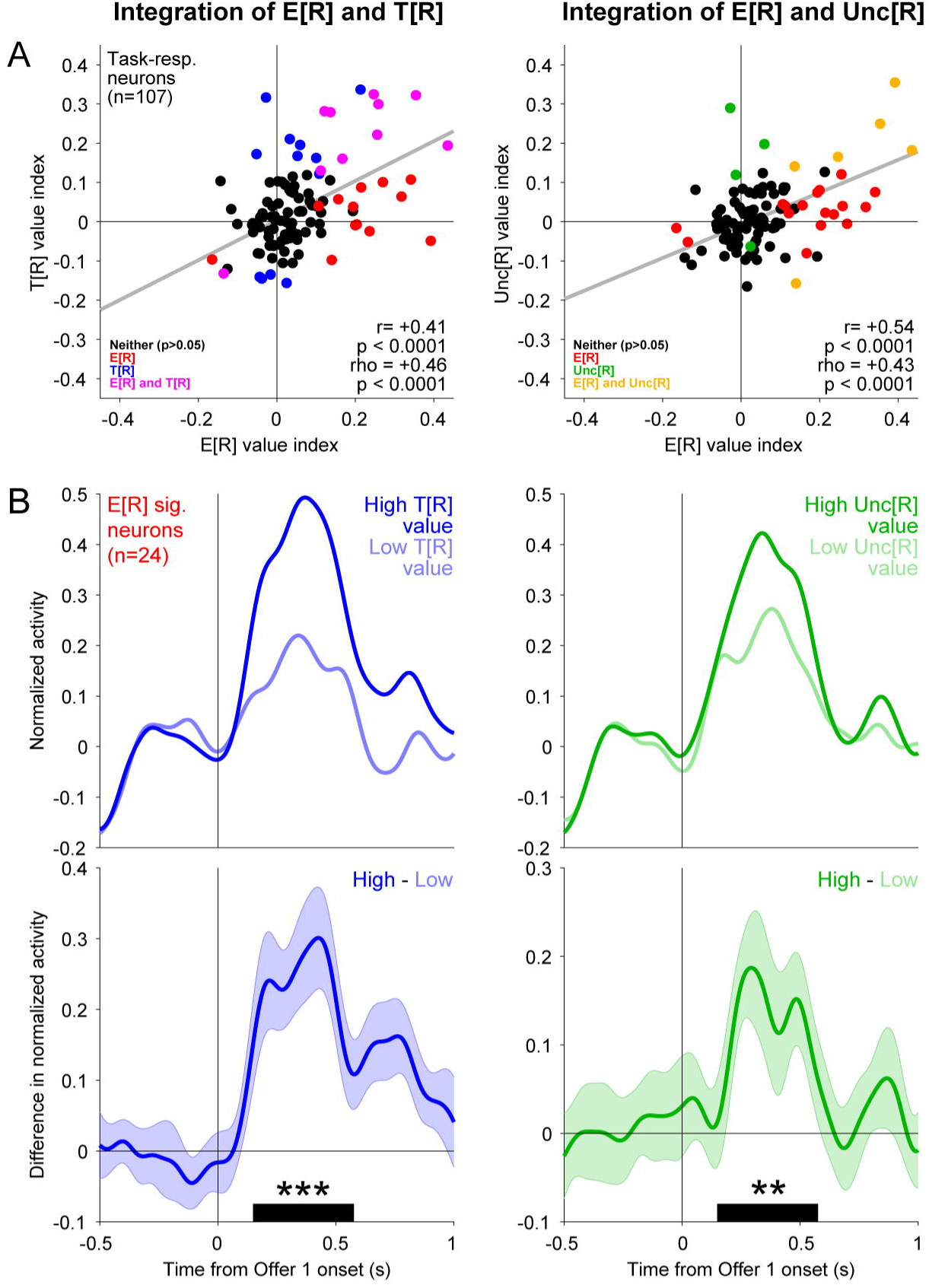
DRN expected reward signals integrate the value of delay or uncertainty. (A) Scatter plots showing E[R] vs T[R] (left) or E[R] vs Unc[R] (right) value signals across task-responsive neurons. E[R] signals are strongly positively correlated with T[R] and Unc[R] (Pearson and Spearman correlations, two-tailed permutation tests). Gray lines show Type 2 regression. One outlier with an extreme E[R] value index was omitted from the scatters for presentation purposes (E[R] value index = 1.0, sig.; T[R] value index = 0.14, n.s.; Unc[R] value index = 0.30, sig.). Colors indicate that neither (black), the x-coordinate (red), the y-coordinate (blue), or both (magenta) are significant (p < 0.05). (B) Top: Average normalized activity for E[R]-signaling neurons on trials where Offer 1 T[R] (left) or Unc[R] (right) was valued in the direction of the neuron’s E[R] value signal (Higher) or the opposite (Lower) (see Methods). Bottom: Difference in the activity shown in B. Error bars denote s.e.m. Average discrimination during the offer response analysis window (150-575 ms after Offer 1 presentation, black bar) was significant (two-tailed signed-rank test, ** p<0.001, *** p<0.001).

### DRN neurons integrate cognitive and physical rewards

So far, we have demonstrated that DRN neurons can signal the value of offers, integrating the expected amount, delay, and uncertainty of juice, a physical reward. These data so far are consistent with the notion that DRN neurons can signal the overall value of an offer. However, if DRN neurons signal overall value, integrating all motivational variables, then they should also signal the value of non-physical or cognitive rewards.

To test this, we took advantage of recent findings showing that monkeys, like humans, often place high value on obtaining advance information to resolve uncertainty about upcoming reward outcomes^42,45,52–56^. Importantly, they value this information even when they cannot use it to influence the physical reward outcome, suggesting that they value resolving uncertainty even when this has no objective, instrumental value for obtaining physical rewards. To measure DRN neurons’ responses to this kind of uncertainty-resolving information, we included two types of offers: informative (‘Info’) and non-informative (‘Noinfo’) (Figure 6A). After monkeys chose an Info offer, a cue would appear early in advance of reward delivery that highlighted the outcome bar, thus resolving reward uncertainty (Figure 6A; Info, ‘Cue’). Conversely, after monkeys chose a Noinfo offer, a visually matched control cue would appear that indicated a random bar, thus leaving reward uncertainty unresolved until just before reward delivery, when the upcoming outcome was revealed (Figure 6A; Noinfo, ‘Reveal’). Consistent with previous work showing that the value of information scales strongly with reward uncertainty^42,45,57,58^, both monkeys significantly scaled up the value of information with increasing Unc[R] (Figure 6B).

**Figure 6.**
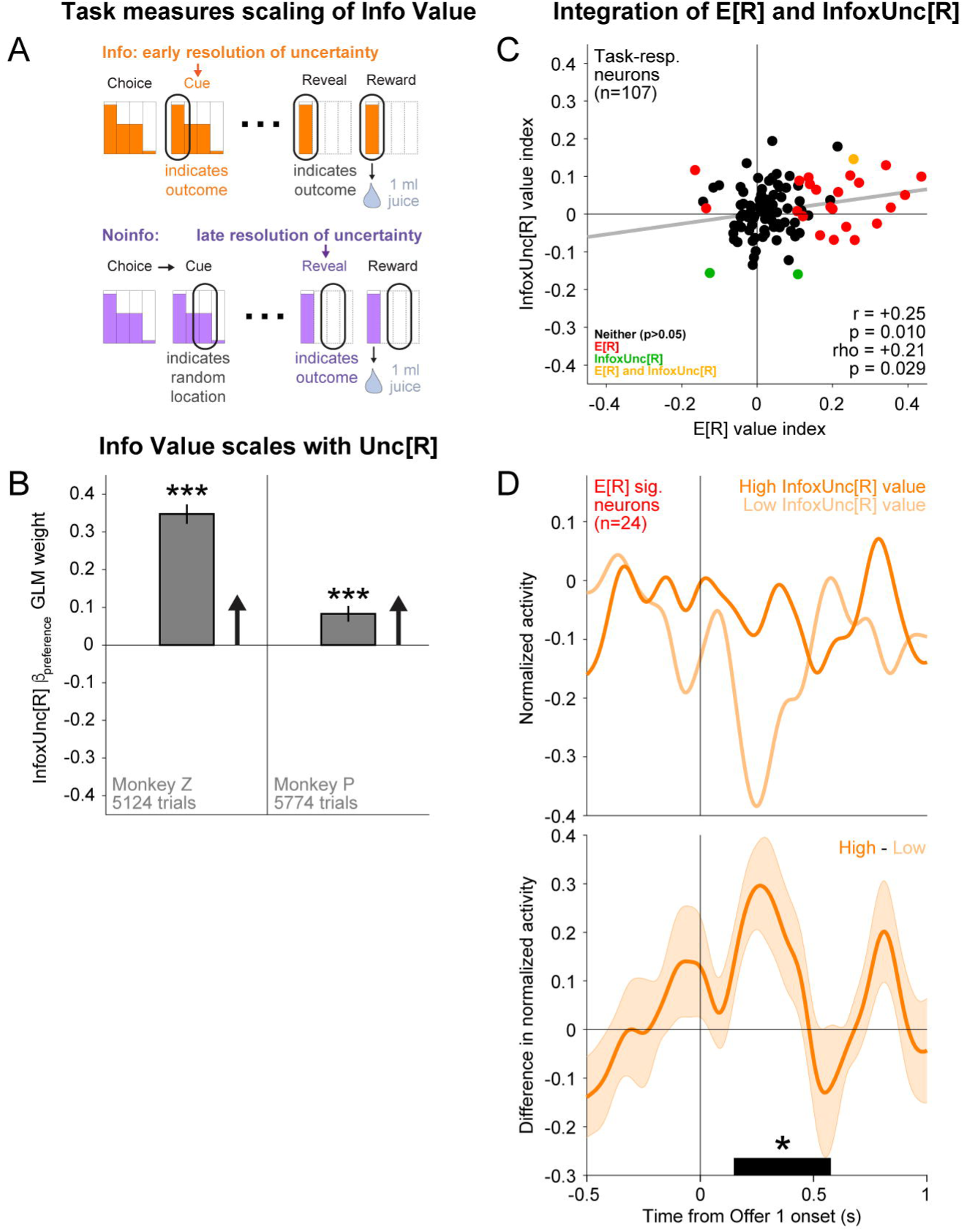
DRN expected reward signals integrate the value of information to reduce uncertainty. (A) The task also measured the value of information to reduce uncertainty about upcoming rewards. Trials could either be informative (Info, orange), for which reward uncertainty would be resolved soon after choice, or non-informative (Noinfo, purple), for which reward uncertainty would not be resolved until immediately before reward delivery (see Methods). One outlier with an extreme E[R] value index was omitted from the scatter for presentation purposes (E[R] value index = 1.0, sig.; InfoxUnc[R] value index = 0.10, n.s.). (B) Bar plot showing fitted InfoxUnc[R] interaction weight (see Methods). Error bars denote s.e.m. Monkeys strongly scaled the value of information with uncertainty. (C) Same as Figure 5A, but for E[R] vs InfoxUnc[R] value signals. E[R] and InfoxUnc[R] are strongly positively correlated (Pearson and Spearman correlations, two-tailed permutation tests). Gray line shows Type 2 regression. Colors indicate that neither (black), the x-coordinate (red), the y- coordinate (blue), or both (magenta) are significant (p < 0.05). (D) Same as Figure 5B but showing differences in normalized activity between Info and Noinfo, splitting trials based on Offer 1 Unc[R].

Having replicated previous findings that monkeys integrate the values of juice reward and uncertainty-resolving information to make their decisions, we next asked whether DRN neurons integrate the values of these cognitive and physical rewards in a similar manner. To quantify neurons’ signals for the value of information to reduce uncertainty, we computed an InfoxUnc[R] value index based on the fitted interaction of Info and Unc[R], analogous to the attribute value indices for physical reward (see Methods). Consistent with integration of cognitive and physical rewards, InfoxUnc[R] and E[R] value indices were correlated across task-responsive neurons (Figure 6C). Moreover, on average, E[R]-signaling neurons signaled InfoxUnc[R] in the same direction as E[R] and with phasic dynamics that resembled how they signaled E[R] (Figure 6D). Thus, DRN neurons can integrate the values of cognitive and physical rewards.

Finally, to estimate how strongly neurons’ offer responses code an offer’s overall value, integrating across its many attributes, we computed a value coding index for each neuron using a recently-employed approach^42^ (see Methods). As suggested by the strong pairwise patterns of integration detailed above, we saw that DRN neurons tended to have very high value coding indices, often very close to the maximum index, consistent with coding of an offer’s overall value, integrating across its many attributes (Supplementary Figure 2).

### DRN neurons track changes in value after decision-making by signaling reward prediction errors

Finally, DRN is thought to continuously track reward states on multiple timescales, potentially containing neurons sensitive to reward predictions or changes in reward predictions^2,18,27–29,31,33,35,38,40,41^. This raises the possibility that the DRN neurons we found to signal value predictions based on offers may continue to track changes in value predictions based on post-decision feedback. We thus asked whether and how DRN neurons process post-decision feedback.

We first characterized DRN neurons’ responses to post-decision feedback, which was provided by the first post-decision event on each trial to indicate the amount of reward to be delivered (Cue on Info trials; Reveal on Noinfo trials; Figure 6A). To do this, we took advantage of several features of our task. Our task precisely manipulated reward predictions and outcomes by offering a diverse set of explicitly communicated reward distributions to the monkeys. This let us precisely test how responses to post-decision feedback were related to (a) the animal’s initial prediction about the offer’s reward value (i.e. E[R] of the chosen offer), before viewing the feedback, (b) the animal’s updated prediction of the offer’s reward value, after viewing the feedback and finding out the actual outcome to be delivered, and (c) the difference between the updated and initial predictions, known as reward prediction error (RPE), which is thought to guide learning and motivation^59–62^. In addition, our task let us test how consistently neurons encode these variables. Specifically, we can test whether neurons encode these variables consistently for feedback with different sensory features and timing^63–66^ (Cue on Info trials vs Reveal on Noinfo trials), consistently across their full range of values (for example, do neurons respond when the actual outcome is better than predicted, worse than predicted, or both)^67,68^, and in a signed or unsigned manner^28,66,69^.

We found that many DRN feedback responses strongly resembled a positively signed RPE signal. For example, consider the neuron in Figure 7A. This neuron was excited when feedback indicated that the reward outcome would be better than predicted (positive RPE), inhibited when feedback indicated that the reward outcome would be worse than predicted (negative RPE), and had little change in activity when feedback indicated that the reward outcome would be the same as predicted (zero RPE). The neuron did this consistently across both task events that provided post-decision feedback, the Cue for Info offers and the Reveal for Noinfo offers (Figure 7A).

**Figure 7.**
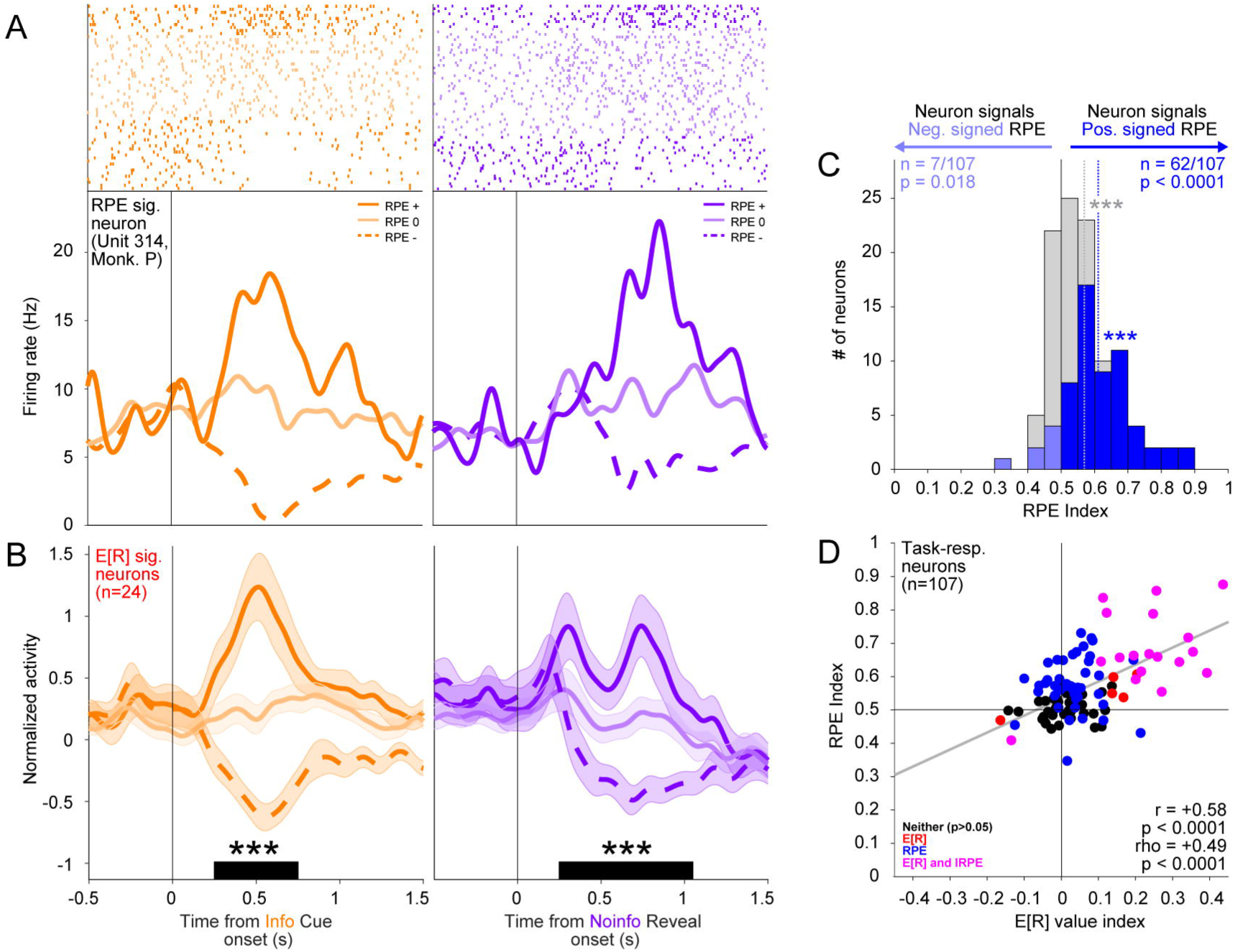
DRN neurons that signal expected reward during decision-making signal reward prediction errors after decision-making. (A) Rasters and SDFs showing an example neuron’s responses to Info Cue (left, orange) or Noinfo Reveal (right, purple) on trials where it elicited an RPE that was positive (RPE +, solid, dark), 0 (RPE 0, solid, light) or negative (RPE -, dashed, dark). (B) Same as A but showing average normalized firing across E[R]-signaling neurons. Info and Noinfo RPE indices (see Methods) were significant (two-tailed signed-rank test, *** p<0.001). Activity was analyzed in the analysis windows shown in the black bars (250 to 750 ms post-cue, Info; 250 to 1000 ms post-reveal, Noinfo). (C) Histogram showing RPE indices (see Methods) of task-responsive neurons. Signals are predominantly positive (dashed lines show mean; median significantly different from 0 by signed-rank test). The proportion of neurons with significant positive RPE indices is large and strongly significant, while the opposite is small and weakly significant (two-tailed binomial tests). (D) Scatter plot showing RPE indices vs E[R] value signals. RPE indices and E[R] value signals were strongly positively correlated (Pearson and Spearman correlations, two-tailed permutation tests). Gray line shows Type 2 regression. One outlier with an extreme E[R] value index was omitted from the scatter for presentation purposes (E[R] value index = 1.0, sig.; RPE index = 0.84, sig.) Colors indicate that neither (black), the x-coordinate (red), the y-coordinate (blue), or both (magenta) are significant (p < 0.05).

To test whether DRN feedback responses had the key properties needed to function as a positively signed RPE signal, we quantified how each neuron’s responses discriminated between positive, zero, and negative RPEs (based on ROC analysis; Methods). First, we found that DRN feedback responses consistently and bidirectionally signaled both positive and negative RPEs. That is, they were more active when reward feedback was better than predicted and were less active when reward feedback was worse than predicted. To test this, we computed separate RPE indices for how neurons discriminated positive RPE from zero RPE (+RPE index) and zero RPE from negative RPE (-RPE index; Methods). As expected for a positively signed RPE signal, these two indices were strongly positively correlated (Supplementary Figure 3A). Second, we found that DRN feedback responses consistently signaled RPEs for both distinct feedback-related events in the task. To test this, we computed separate RPE indices for responses to the Info offer Cue (Info RPE index) and the Noinfo offer Reveal (Noinfo RPE index; Methods). As expected for a positively signed RPE signal, these two indices were strongly positively correlated (Supplementary Figure 3B). Third, we found that DRN feedback responses commonly responded in opposite directions to the initial reward prediction (the offer’s expected value) and the updated reward prediction (the actual reward outcome that would be delivered), consistent with signaling the difference between them. To test this, we fit each neuron’s activity using a GLM with separate terms for predicted and delivered reward. As expected for a positively signed RPE signal, neurons predominantly signaled delivered amounts positively and predicted amounts negatively (Supplementary Figure 4A). In sum, this activity profile is consistent with a positively signed RPE signal.

For each neuron, we summarized its RPE signal using a single RPE index computed as the mean of its + and - RPE indices for Info and Noinfo trials. The mean RPE index across all task-responsive neurons was strongly and significantly positive (+0.57, p<1E-16; two-tailed signed-rank test), consistent with a positively signed RPE signal like that of the example neuron. A large proportion of neurons had significant RPE indices (n=69/107, 64%), which were overwhelmingly positive. The proportion of neurons with significantly positive RPE indices was much greater than chance (n=62/107, p<1E-16; two-tailed binomial test). Interestingly, there was also a significant, albeit small, proportion of neurons with significantly negative RPE indices (n=7/107, p=0.018; two-tailed binomial test), in line with prior observations of some DRN neurons with negative reward discrimination^27,34–36,70^. There was also a subpopulation of neurons with feedback responses that were less influenced by reward expectation (Supplementary Figure 4A-C), again in line with prior reports^29,33,34^.

We then examined the relationship between offer value signals during decision-making and RPE signals after decision-making. E[R] value indices and RPE indices were strongly correlated across task-responsive neurons (Figure 7D). Similarly, signed value coding indices and RPE indices were also strongly correlated across strongly attribute-responsive neurons (Supplementary Figure 5). Moreover, E[R]-signaling neurons signaled RPE in the same direction as Offer E[R] and with similar phasic dynamics (Figure 7B), and we observed more neurons than expected by chance that had significant RPE and E[R] value indices in the same direction (n=18/107, p<1E-16; two-tailed binomial test). E[R]-signaling neurons were also significantly more likely to respond to post-decision feedback by both positively signaling delivered amounts and negatively signaling predicted amounts, as expected for a positively signed RPE signal (Supplementary Figure 4C,D).

In summary, we find that DRN neurons that signal the value of offers during decision-making also commonly signal positively signed, bidirectional RPEs in response to feedback after decision-making. Thus, DRN signaling of offer attributes during decision-making may reflect a more generalized role in tracking changes in value predictions across task epochs and behavioral contexts.

## Discussion

We found that DRN neurons signal expected reward amount, reward delay, and reward uncertainty during decision-making, in a manner reflecting the integrated value that guides monkeys’ choices. Additionally, we demonstrate that DRN neurons signal RPEs in response to post-decision feedback, suggesting a broader role in tracking value across task epochs and contexts.

Value is a complex construct thought to reflect preferences over many distinct attributes and to regulate value-based decision-making and other value-guided behaviors^71–73^. Our results provide the first evidence that DRN reflects the integrated value of offers during decision-making based on multiple attributes of juice reward, including expected amount, delay, and uncertainty, as well as the value of uncertainty-resolving information. There are many other factors that influence value and which other brain areas have been shown to process during value-based decision-making^43,74–87^, such as effort or punishment. While we did not assess all of these factors, our findings that DRN signals integrated value based on multiple attributes of physical and cognitive rewards, in a systematic manner across the neural population, raise the possibility that they can integrate an even wider range of the many motivational variables that govern decisions.

We found that DRN neurons can track changes in value predictions by signaling positively signed, bidirectional RPEs. This was somewhat unexpected because some previous results reported that DRN neurons may primarily signal the value of outcomes themselves^29,33,34^, or respond positively to both positive and negative RPEs, as if signaling surprise^28^. However, other results have been consistent with positively signed RPE, with DRN neurons responding more strongly to unexpected vs expected rewards^29,36^ and cues predicting more vs less reward^28,29,34,36^. It is possible that different task conditions and behavioral strategies engender different response patterns in DRN neurons. In addition, differences in recording locations could target DRN subregions with distinct functions^88,89^. Indeed, recent studies suggest that axons originating from more caudal subregions of DRN are predominantly inhibited by rewards^90^, whereas neurons in more rostral subregions, potentially more analogous to our recording locations, are predominantly excited^91^. There is likely also heterogeneity within subregions of DRN; for example, we also observed a subset of neurons whose responses to post-decision feedback primarily reflected the reward amount to be delivered, with relatively weak modulation by the predicted amount (Supplementary Figure 4A-C). Finally, it is also possible that some aspects of DRN function differ across species, given the known differences in DRN nuclear organization and projection patterns between rodents and primates^92^.

DRN receives input from several areas that could support value and RPE computations, including regions of prefrontal cortex (PFC), lateral hypothalamus, ventral pallidum (VP), and the lateral habenula (LHb)^92–95^. PFC areas involved in emotion and cognition project strongly to DRN, including the anterior cingulate cortex (ACC) and dorsolateral PFC (DLPFC)^96,97^. ACC is known to signal multiple motivational variables^77,98–102^, and we recently showed that VP signals partially integrated offer attributes in the same task used here^42^. Also, both ACC and VP have been implicated in processing motivational predictions and outcomes^99,103^. DRN could utilize these inputs from ACC and VP in flexible ways depending on changes in internal needs such as hunger or thirst, potentially modulated by lateral hypothalamus input^104^, or differing task demands or contingencies, potentially signaled by DLPFC^105^. Among DRN inputs, LHb is notable because it is well-known to potently signal negatively signed RPEs^106,107^. Moreover, we recently showed that it also negatively signals integrated value in this same multi-attribute decision-making task paradigm^42^. LHb has reciprocal connections with DRN and an established role in controlling other neuromodulators^108^. It also receives input from many DRN-projecting areas, including the medial PFC, VP, and lateral hypothalamus^109^. Future studies should investigate the relationship between integrated value and RPE signals and the dynamics of their co-emergence in the DRN-LHb circuit.

DRN is the largest component of the serotonin (5-HT) system, sending projections throughout the brain^110^. In addition to its 5-HT neurons, DRN contains other cell types^111,112^, which project to cortical and subcortical targets and regulate DRN 5-HT neurons^113–115^. While we recorded in areas enriched with 5-HT neurons, where some estimate >60% of neurons release 5-HT^50^, we cannot confirm cell types in our data and it is unlikely that we exclusively recorded 5-HT neurons. DRN also contains glutamate-^113,116,117^, GABA-^113,118^, and dopamine-releasing^70,119–121^ subpopulations that are involved in motivational processing, potentially through projections to downstream areas or local influence over DRN 5-HT neurons^115,118,122–125^. Future studies should examine cell-type heterogeneity in primates during decision-making and other complex behaviors. Nonetheless, dissecting how value is processed within DRN is crucial for understanding the complex role of the 5-HT system in coordinating value-guided computations throughout the brain.

An influential theory of 5-HT function posits an opponent role to dopamine (DA), with phasic modulations signaling punishment prediction errors^2,126^. In contrast, we find that many DRN neurons positively signal offer value and RPEs, which strikingly resembles the responses of midbrain DA neurons in the ventral tegmental area (VTA) and substantia nigra pars compacta (SNc)^60–62^. This is broadly consistent with previous observations of DRN neurons that code reward-related variables positively – some reporting a mix of positive and negative coding neurons^27,34–36,70^ and others reporting predominantly positive^29,30,117,127,128^ – and with recent findings that DRN may contribute to VTA RPE signals through monosynaptic projections onto DA neurons^129^. While these results do not rule out the possibility that certain subpopulations of neurons in subregions of DRN^91^ or other 5-HT nuclei^130^ may indeed play an opponent role to DA, they suggest a new, potentially cooperative relationship between the 5-HT and DA systems^131^.

The cortex receives relatively light DA input compared to striatum^132–134^ and within the frontal lobe, DA projections follow an increasing rostro-caudal gradient, with motor cortex receiving the most DA input and PFC receiving less^135–139^. Thus, other neuromodulators may also be required to supply value-related signals to PFC to support its many roles in value-related processes^140^. 5-HT projections, while widespread, display a pattern of regional distribution that differs from and may complement that of DA. 5-HT projections are more balanced between cortex and striatum^132,133^ and within the frontal lobe, 5-HT projects more strongly to PFC than DA and only lightly to motor areas^136,137,141^. Thus, the DRN 5-HT system could broadcast value-related information to regulate networks at least partially distinctly from DA. The functional significance of these observations requires further investigation.

Our findings that DRN signals integrated value and RPE are broadly in line with existing theories of DRN and 5-HT in signaling reward variables such as ‘beneficialness’^11,37^, ‘the availability of time and resources’^10^, and ‘global reward state’ (long-term average reward rate)^18,40,41^. While these value-related constructs likely cannot fully explain the functions of the entire DRN and 5-HT system, a DRN signal tracking integrated value could explain how the diverse set of DRN projections^89,91,110,111,142^ and complex repertoire of 5-HT receptors^143–148^ coordinately regulate multiple computational and behavioral processes^4,8,10,11,38,126,149–153^. Firstly, DRN and the 5-HT system are theorized to regulate value-based decision-making^3–5,20^. Signaling the integrated value of offers or opportunities as they appear in the environment could support dynamic, online decision-making, potentially through DRN projections to orbitofrontal cortex (OFC)^73,87,154,155^ or LHb^42,156,157^, whose value-related activity is causally involved in decision-making, or DLPFC, which has been implicated in decision processes more generally^80,155,158–160^. Secondly, an early theory of 5-HT proposed a role in mediating behavioral inhibition^161^. While this is often considered in the context of avoiding punishments^4,126,162^, obtaining maximal rewards can also require patience, persistence, and deliberation^10,163–168^, which an integrated value signal could help to regulate^10,169^. Indeed, optogenetic stimulation of DRN 5-HT neurons^170,171^ or their projections to OFC and a region of rodent medial PFC thought to be homologous to ACC^172^ promotes waiting for rewards^173^, whereas chemical inhibition of DRN 5-HT neurons^174^ or blockade of 5-HT_1A_ receptors^175^ reduces waiting for rewards. Thirdly, the 5-HT system is theorized to regulate learning and cognitive flexibility^28,150,176–192^. Integrated value signals could indicate that time and resources are available for safe and fruitful learning^10,28,176,183,190,193^, by promoting exploration and information-seeking through DRN projections to ACC and ventrolateral PFC^99,194,195^ or by accelerating the rate of learning through 5-HT-mediated neuroplasticity across the brain^196^. Additionally, DRN’s RPE signal could update value or policy estimates, directly through 5-HT-mediated long-term depression^197,198^ or indirectly by influencing DA-mediated long-term potentiation^197,198^ through DRN projections to VTA DA neurons^129,199^.

Major depressive disorder (MDD) is perhaps the psychiatric illness most often linked to DRN and 5-HT system dysfunction^200,201^, and drugs targeting 5-HT transmission remain the main pharmaceutical therapeutics^202^, including conventional drugs such as selective serotonin reuptake inhibitors (SSRIs) and emerging ones such as psilocybin, a serotonergic psychedelic^203–205^. However, computational frameworks for understanding MDD are limited, in part due to its poorly defined neural correlates and complex, heterogenous clinical presentations^206–208^. MDD is characterized by wide-ranging symptoms, mainly low mood and anhedonia, but also other deficits such as psychomotor slowing, sleep abnormalities, and impaired cognitive function^209^. An integrated value signal that modulates multiple computational and behavioral processes could also contribute to the wide-ranging symptomatology of MDD and link it to DRN and the 5-HT system. A signal tracking the environment’s changes in integrated value could modulate mood, which has been hypothesized to represent ‘environmental momentum’^210–213^. Similarly, lower estimates of value could contribute to anhedonia, defined as loss of interest or pleasure from objects or activities one previously found rewarding^207,214^, and negativity biases^215^. In turn, mood and value could regulate various behavioral parameters, including action vigor^207,216^, sleep^207,217^, and decision-making^206,207,218^. Moreover, while theories of serotonergic antidepressant mechanisms have been proposed at the cellular and psychological levels, it is difficult to unify them without a computational understanding of DRN and the 5-HT system. The therapeutic efficacy of both SSRIs and psychedelics has been linked to their effects on the activity of DRN neurons, 5-HT receptor availability, and neuroplasticity^219–224^, which may then manifest as adaptive psychological changes, such as reduced negativity biases and improved cognitive flexibility^219,225–229^. The efficacy of these drugs in treating other psychiatric illnesses such as anxiety or obsessive-compulsive disorder may also depend on similar mechanisms^230–237^. Thus, an integrated value signal that is processed within DRN could link these neurobiological and psychological effects transdiagnostically. In summary, in addition to guiding theories of the 5-HT system in adaptive behavior, our findings about the neural computations of DRN neurons will also inform our understanding of the neural and computational basis of psychiatric disease, potentially leading to improved therapeutic avenues.

## Methods

### Experimental model and subject details

Two adult male monkeys (Macaca mulatta; Monkey Z and P; ages: 7-10 years old) were used for experiments. All of these procedures conform to the Guide for the Care and Use of Laboratory Animals and were approved by the Institutional Animal Care and Use Committee at Washington University.

### Data acquisition

A plastic head holder and plastic recording chamber were fixed to the skull under general anesthesia and sterile surgical conditions. The chambers were tilted laterally (28° in Monkey Z; −34° in Monkey P) and aimed at DRN. After the monkeys recovered from surgery, they participated in behavioral and neurophysiological experiments.

While the monkeys participated in the behavioral procedure, we recorded single neurons in DRN. Electrode trajectories were determined with a 1 mm-spacing grid system and with the aid of MR images (3T) obtained along the direction of the recording chamber. This MRI-based estimation of neuron recording locations was aided by custom-built software^238^. In addition, in order to further verify the location of recording sites, after a subset of experiments the electrode was temporarily fixed in place at the recording site and the electrode tip’s location in the target area was verified by MRI (Figure 2A).

Electrophysiological recordings were performed using multi-contact electrodes (Plexon V-probes, 32 or 64 channels, 50 μm spacing) inserted through a stainless-steel guide tube and advanced by an oil-driven micromanipulator (MO-97A, Narishige). Signal acquisition (including amplification and filtering) was performed using an OmniPlex 40 kHz recording system (Plexon). Spike sorting was performed offline using publicly available software (Kilosort2) to extract clusters from the recordings, followed by manual curation to identify sets of clusters that corresponded to single neurons, the spans of time when they were well isolated, and whether they were in the regions of interest. DRN was identified based on neurons’ electrophysiological characteristics, firing patterns, and locations relative to anatomical landmarks estimated from MRIs and recordings at multiple grid locations. Specifically, we relied on visuo-auditory responses in the superior colliculus, distinct electrophysiological characteristics in the periaqueductal gray, oculomotor responses in the trochlear nucleus, and axonal characteristics in the medial longitudinal fasciculus. Neuronal and behavioral analyses were conducted offline in Matlab (Mathworks, Natick, MA).

Eye position was obtained with an infrared video camera (Eyelink, SR Research). Behavioral events and visual stimuli were controlled by Matlab (Mathworks, Natick, MA) with Psychophysics Toolbox extensions. Juice, used as reward, was delivered with a solenoid delivery reward system (CRIST Instruments).

### Behavioral task

Each trial began with the appearance of a white fixation point at the center of the screen which the animal was required to fixate with its gaze. The task proceeded once the animal maintained fixation continuously for 0.25 s and at least 0.5 s had passed since fixation point onset. If the animal did not fixate within 5 s of fixation point appearance, or broke fixation before 0.5 s had passed since fixation point onset, the trial was counted as an error, moved immediately to the inter-trial interval (ITI), and repeated until it was performed successfully. After the fixation requirement was successfully completed, the fixation point disappeared and there was no longer any gaze requirement to advance the task. Simultaneously with fixation point disappearance, the first offer appeared on the screen. After 1 s, the second offer appeared on the screen. After a further 0.5 s, the choice period began. An offer was counted as chosen once the animal held its gaze on it continuously during the choice period for a fixed duration (0.4 s in Monkey Z; 0.5 s in Monkey P). For Monkey Z, if the monkey did not choose an offer within 5 s of the start of the choice period, the computer randomly selected an offer and the trial proceeded as if the animal had chosen it. For Monkey P, if the monkey did not choose an offer within 5 s of the start of the choice period, the current trial would end and a repeat trial would begin after the normal ITI, on which the same offers would be shown. For both monkeys, failure to choose within 5s occurred rarely, and computer-chosen trials and repeat trials as well as error trials were excluded from analysis unless otherwise noted. The size of each offer stimulus was approximately 6 x 5 degrees of visual angle. The locations of the two offers on the screen were randomly selected on each trial from a set of three possible locations (center, 10 degrees left of center, 10 degrees right of center).

Each offer featured a reward distribution that was visually depicted with 4 bars, with the height of each bar proportional to the volume of its juice reward, up to a maximum size Rmax (1.2 mL) corresponding to the maximum bar height. If an offer was chosen, one of its 4 bars was selected uniformly at random as the outcome, and hence the juice volume indicated by that bar was delivered to the animal at the time of outcome delivery. Juice was delivered through a metal spout placed directly in the mouth, to ensure that anticipatory actions such as licking were not required to obtain the reward and did not influence the amount of reward that was received.

We presented offers with multiple types of reward distributions. In brief, when randomly generating each offer on each trial, the offer first had its expected reward selected uniformly at random from a pre-specified set of possible expected rewards. To reduce the number of trials with trivial decisions where one offer was much better than the other, Offer 2’s expected reward was constrained to be within a pre-specified range of Offer 1’s expected reward. Then, the offer had its reward distribution randomly drawn from one of the following types: Safe distributions with 100% probability of delivering a specific reward amount; 25/50/25 distributions with 25, 50, and 25% chances of delivering small, medium, and large reward amounts, respectively, where the medium amount was equal to the mean of the small and large amounts; and 50/50 distributions with 50% chances of delivering small or large amounts. Then, after the offer’s distribution type was selected, its reward range was randomly drawn from a pre-specified set of ranges (50% or 70% of Rmax).

Each offer also could be informative (Info) or non-informative (Noinfo), indicated by its bars having distinct visual textures (we used complex visual textures sourced from images of scenes; for clarity of presentation, these are depicted in the figures as simple colors, i.e. orange or purple). If an informative offer was chosen, a visual cue was later presented that highlighted the single bar that was selected to be the outcome on that trial. If a non-informative offer was chosen, the same visual cue was presented at the same time but highlighting a single random bar. Thus, choosing an informative offer gave early access to information about the trial’s upcoming reward outcome. For monkey P, to reduce choice complexity, informativeness was always matched between Offer 1 and 2.

After the choice was made, the unchosen offer disappeared, and then a sequence of timed events occurred. First, at time Tcue, a visual “cue” appeared on the chosen offer in the form of a black box highlighting one of its bars. Then, at time Treveal, a “reveal” occurred in which three of the bars disappeared and only one bar remained, and then a further 1.1 s the reward was delivered (thus, Treveal fully determined the reward delay, with T[R] = Treveal + 1.1 s). The remaining bar always indicated the trial’s outcome, and hence, always provided full information about the trial’s upcoming outcome. This was to ensure that both informative and non-informative offers did eventually provide complete information about the trial’s outcome before it was delivered, to ensure that animals always had adequate time to physically prepare to consume the reward. Finally, the offer disappeared 8 s after the choice was made, and then a 1.2 s ITI occurred before the next trial began.

Each offer’s event timing, including Tcue and Treveal, was visually depicted as a large “clock” in the form of a horizontal bar placed at the bottom of the offer. This has been described in full detail previously^42^. The horizontal extent of the clock represented the full-time duration between choice and the end of the trial (8 s). The clock was divided into three sequential segments, representing the segments of time (A) from the choice to the cue (Tcue), (B) from the cue to the reveal (Tadvance = Treveal - Tcue), and (C) from the reveal to the end of the trial (8 s - Treveal). The time of reward delivery was not explicitly indicated but was always exactly 1.1 s after the reveal. The three segments of the clock each had distinct visual textures, which were also distinct for informative vs. non-informative offers (again, for clarity of presentation, these are depicted in figures as simple colors). In addition, two thick black vertical lines indicated the time of the cue and the time of the reveal. Finally, to help animals learn how the clock corresponded to real world time, and to help them anticipate the time of each event during each trial, we animated the clock. Specifically, after the choice was made, a “hand of the clock” appeared at the left edge of the clock, in the form of a thick black rectangle whose position on the clock indicated the current time during the trial. The hand moved from left to right at a constant speed as the trial proceeded. When the hand touched the first black vertical line indicating Tcue, an animation played lasting 0.6 s in which the vertical line moved upward to the center of the cued bar and then expanded to become the cue. When the hand touched the second black vertical line indicating Treveal, an animation played lasting 0.6 s in which the vertical line moved upward to the center of the chosen offer and then became an expanding rectangular region with the background color of the screen, “erasing” three of the bars thus leaving only one bar remaining. Finally, when the hand touched the right edge of the clock, the trial ended.

We presented offers with different cue and reveal (and thus, reward delivery) timing. For monkey Z, reveal timing was drawn jointly with cue timing. Specifically, the pair of event times, (Tcue, Treveal), was drawn uniformly at random from the set of all possible such pairs meeting the following requirements: Tcue was between 0.4-5.8 s; Treveal was between 1.2-6.7 s; Tcue came at least 0.8 s before Treveal; Tcue and Treveal were both multiples of 0.1 s. For monkey P, Treveal was drawn uniformly at random from the set (2.0, 2.9, 5.6, 6.5 s), while Tcue was fixed at 0.5 s to maximize the value of information^42^ and reduce choice complexity.

### Data analysis

All statistical tests were two-tailed unless otherwise noted. All statistical significance corresponds to p < 0.05 or a 95% confidence interval excluding 0 unless otherwise noted. Neural firing rates in response to each offer were analyzed in a time window 150-575 ms after offer onset. Firing rates for each neuron were converted to normalized firing rates by z-scoring. Specifically, for each neuron, we computed a vector of firing rates representing its full activity throughout the task by binning its data from all times during all trials in non-overlapping 500 ms bins. We then normalized the neuron’s firing rates in our analyses by subtracting the mean of that vector and dividing by the standard deviation of that vector. For plotting purposes, activity was smoothed using a Gaussian kernel (σ = 50 ms).

#### Task-responsiveness

Significant task-responsiveness was defined as variance in neural activity across all task events. To determine whether recorded neurons had significant task event-related modulations, we computed P values by comparing activities across different trial or types within the time windows of: −800 to 0 ms from Cue or Reveal onset and 0 to 900 ms from Offer 1, Offer 2, Cue, Reveal, or Reward delivery onset (Kruskal–Wallis test; PL<L0.05). The windows were chosen such that they included key neuronal modulations and were wide enough to avoid bias towards a particular response pattern (e.g., phasic versus tonic). For Post-Offer 1, Post-Offer 2, Pre-Cue, and Pre-Reveal windows, 1-way comparisons were made for E[R], Unc[R], T[R], and Info, split into quantiles with 5, 5, 4, or 2 (Info vs Noinfo) levels, respectively. For Post-Cue and Post-Reveal windows, the same 1-way comparisons were made, but with the addition of delivered reward amount minus chosen offer E[R], split into quintiles. For the Post-Reward delivery window, a 1-way comparison was made only for delivered reward amount, split into quintiles. The P values were then combined using Fisher’s combined probability test^239,240^. This nonparametric method has fewer assumptions than parametric methods.

#### Offer attribute models for behavior and neuronal activity

We fit each individual’s binary choice data using a generalized linear model (GLM) designed to model a standard decision-making formulation in which values are computed for each of the two offers, and then the resulting choice probability is a logistic function of the difference in the values:

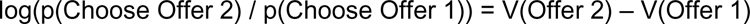

And the value of each offer i is a linear weighted combination of the offer’s vector of n attributes <x_i,1_,x_i,2_,x_i,3_,…,x_i,n_>:

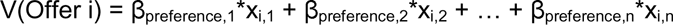

Thus, the resulting model was a GLM for binomial data with a logistic link function, with the equation: log(p(Choose Offer 2) / p(Choose Offer 1)) =

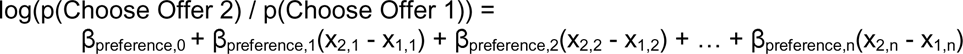

…where β_preference,0_ is the subject’s choice bias.

For analysis of neuronal data, we used an analogous GLM to fit neuronal firing rates in response to Offer 1, for normal data with an identity link function (i.e. equivalent to ordinary linear regression):

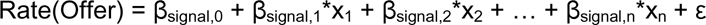

…where β_0_ is the neuron’s mean or baseline response to the offer and ε is the error term.

There were also several aspects of the model fitting procedure that held across models. First, to permit more interpretable comparisons between attributes, unless otherwise noted, all attributes that were not binary were standardized by z-scoring. Second, in addition to the terms described below related to the values of the offers, all models also included terms for response biases, including the location in space of each offer and the sequential order in which the offers were presented. For both behavioral and neural data, this included binary attributes representing whether or not the offer was on the left side of the screen and whether or not the offer was on the right side of the screen.

For monkey Z’s data, there were many possible variables to consider, since the data included manipulations of both Tcue and Treveal, in addition to all of the other task variables described above. Therefore, we focused on a core set of attributes that most strongly governed the subjective value of offers across monkeys, adapted from the set derived using a model selection procedure in our previous study^42^: E[R], T[R], Unc[R], E[R] x T[R], E[R] x SD[r], Info, Info x E[R], Info x T[R], Info x Unc[r], Info x Tadvance, Tadvance x Unc[R], Info x Tadvance x Unc[R]. For monkey P’s data, the main effect of Info and interactions involving Tadvance were excluded, since Info was always matched between offers and Tadvance was fully determined by T[R]. We then defined a model with these attributes for our analysis of both monkey behavior and neuronal activity in this task.

For each neuron, we computed its attribute value indices simply as the fitted β weight for each attribute, sign-flipped based on the direction of the recorded monkey’s preference for each attribute such that positive attribute value signals indicate higher firing for offers with higher value based on that attribute.

#### Analysis of reward prediction error coding

To quantify each neuron’s coding of conventional juice reward prediction errors, we computed a reward prediction error index (RPE Index) using the following approach. First, we defined the RPE on each correctly performed trial as the difference between the delivered reward amount and expected reward amount (i.e. the chosen offer’s E[R]). Next, we selected the subsets of trials which fell into each of four categories, based on the 2×3 combinations of the chosen offer’s informativeness (Info or Noinfo) and the resulting RPE (+,0,-), corresponding to RPEs > +0.1 mL juice, < +0.1 mL juice and > −0.1 mL juice, or < −0.1 mL juice). We then analyzed single trial firing rates on these trials after two task events where RPEs commonly occurred in our task: a post-cue time window 250-750 ms after cue onset for Info trials, and a post-reveal time window 250- 1050 ms after reveal onset for Noinfo trials. We then used this activity to compute separate measures of the strength of the positive and negative components of the RPE responses separately for Info vs Noinfo trials. Specifically, we first computed 4 separate indices as ROC(X,Y), the ROC area under the receiver operating characteristic curve^241^ between the set of single trial normalized activities in condition X vs. condition Y, such that the ROC area is > 0.5 if condition Y generally has higher activity than condition X, and < 0.5 if condition Y generally has less activity than condition X, and 0.5 if the distributions are the same. Significance was assessed by Wilcoxon the rank-sum test. For each index, X and Y were defined as:

For Info RPE+, X and Y were Info trials where RPE was 0 or +, respectively.
For Info RPE-, X and Y were Info trials where RPE was - or 0, respectively.
For Noinfo RPE+, X and Y were Noinfo trials where RPE was 0 or +,respectively.
For Noinfo RPE-, X and Y were Noinfo trials where RPE was - or 0, respectively.

To assess the relationship between RPEs elicited by Info Cues vs Noinfo Reveals, we computed Info and Noinfo RPE indices by averaging Info RPE+ and Info RPE- or Noinfo RPE+ and Noinfo RPE- indices, respectively. P-values for these averaged indices were computed from the p-values of the two constituent measures using Fisher’s combination test^239,240^. Finally, the total RPE Index was computed as the mean of the Info and Noinfo RPE indices and its p-value was again computed from the p-values of these two constituent measures using Fisher’s combination test. Since RPE is defined as the difference between the delivered reward amount and the predicted reward amount (i.e. the chosen offer’s E[R]), RPEs are positively correlated with the delivered reward and negatively correlated with the predicted reward amount. In order to verify that apparent RPE signaling did not simply reflect signaling of either the delivered reward amount or predicted reward amount, we also computed delivered reward amount and predicted reward amount indices. Specifically, we fit each neuron’s normalized activity to two GLMs for normal data with an identity link function (i.e. equivalent to ordinary linear regression). The models were fit to activity in the post-cue window on Info trials or the post-reveal window on Noinfo trials. Specifically, the models were expressed as:

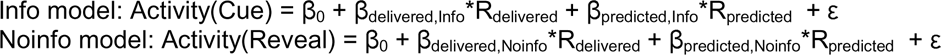

…where β_0_ is the neuron’s mean or baseline response to the Cue or Reveal, R_delivered_ and R_predicted_ are the delivered and predicted reward amounts, and ε is the error term.

The delivered reward amount and predicted reward amount indices were then computed by averaging the corresponding fitted β weights across the Info and Noinfo models, and p-values for these averaged indices were computed from the p-values of the two constituent measures using Fisher’s combination test:

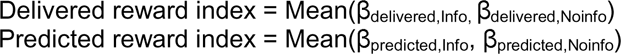

#### Analysis of peri-stimulus activity

To analyze cross-validated timecourses of Offer 1 E[R], T[R], and Unc[R] responses across the population of task-responsive neurons (Figure 3B), we used the following procedure. For each neuron, its data were split into two cross-validation folds, 1 and 2, which included odd-numbered and even-numbered trials, respectively. The neuronal offer attribute model was fit to each fold’s data to obtain β_signal_ weights. For each attribute, we split trials into High vs Low based on Offer 1 (Larger vs Smaller E[R], median split; Shorter vs Longer T[R], median split; Uncertain vs Certain Unc[R]), separately for each fold. We computed the attribute’s effect on the Offer 1 response as the difference in mean normalized firing rate during the Offer 1 analysis window on High vs Low trials. Then, we computed the total cross-validated effect by taking the mean of the effects in the two folds, after multiplying each fold’s effect by the sign of the β_signal,attribute_ weight derived from the other fold’s effect. This provided a cross-validated measure of the total strength of the attribute effect in units of normalized activity, such that a more positive effect indicates a stronger interaction, regardless of whether the underlying effect on firing rate had a positive or negative sign. Specifically, if the neuron had a true effect that could be measured with a consistent sign in both folds, then its total effect would be positive (regardless of whether the sign of the underlying effect reflected a positive or negative influence on firing rate). Conversely, under the null hypothesis that the neuron had no true effect, the total effect would have an expected value of 0, since each fold would be equally likely to be multiplied by either a negative or positive sign. We then tested whether the median total cross-validated effect across the population of neurons was different from 0 (signed-rank test). We also plotted the full timecourse of the mean normalized activity for High vs Low conditions and total cross-validated effect by computing it using the full spike density functions from each cross-validation fold, sign-flipped as in the analysis above (so that the full timecourse of each neuron’s activity in each fold was normalized using a consistent sign).

To analyze time courses of Offer 1 E[R], T[R], Unc[R], and Info x Unc[R] responses across the population of E[R]-signaling neurons, we used the following procedure. We selected E[R]-signaling neurons based on significance of β_signal,E[R]_ from the neuronal offer attribute model. For E[R], T[R], and Unc[R] (Figure 5B), we again split trials into two categories based on Offer 1, but this time into preferred vs unpreferred, based on animals’ behavior. For E[R] and T[R], these were again median splits of Larger vs Smaller E[R] or Shorter vs Longer T[R], respectively. For Unc[R], these were Uncertain vs Certain Unc[R] for monkey Z and Certain vs Uncertain Unc[R] for monkey P. If a neuron had a negative β_signal,E[R]_, its preferred vs unpreferred trials were exchanged. These categories of trials then represented what a value-signaling neuron should signal as High vs Low value for each attribute. For Info x Unc[R] (Figure 6D), we first split trials into 2×2 categories (Info, Noinfo) x (Uncertain, Certain) based on Offer 1. For each neuron, we computed the difference in Offer 1 response on Info vs Noinfo trials, separately for Uncertain vs Certain trials to obtain Uncertain Info vs Certain Info effects, respectively. If a neuron had a negative β_signal,E[R]_, its Uncertain vs Certain effects were exchanged, to give what a value-signaling neuron should signal as High Info value vs Low Info value.

### Value coding index

We used a value coding index we previously defined^42^ to quantify the extent to which the offer responses of each neuron aligned with our behaviorally-derived estimate of how animals weighted those attributes to compute the offer’s overall subjective value. To do this, we first fit each neuron’s Offer 1 responses using the attribute model described above, which allows the neuron to have separate weights for each of the selected attributes described above that influenced the animal’s choices. We then fit responses with a new model called the “value model”, which only had a single attribute, ‘Value’, equal to the estimated value of the offer derived from the behavioral model (using the procedure described above). Thus, the value model represents the hypothesis that the neuron’s attribute-related activity is fully explained by encoding of a single variable, the offer’s overall subjective value; or equivalently, that the neuron’s weighting of the offer’s many attributes to govern its response, is perfectly correlated with the animal’s weighting of the offer’s many attributes to compute subjective value. Finally, we quantified what fraction of the above-chance variance in the neuron’s activity that was explained by the attribute model could also be explained by the value model. To do this, we used a measure based on R-squared (R_2_) to quantify the fraction of the response variance that each model explained. Specifically, to correct for the fact that the attribute model had more parameters than the value model, we computed the shuffle-corrected R-squared (R^2^_corr_), defined as the R_2_ computed on the neuron’s real data minus the mean of the R^2^ computed on 100 shuffled datasets in which the neuronal offer responses were shuffled across trials. Thus, R^2^_corr_ quantified the fraction of variance of neuron’s offer responses that the model explained, above and beyond the chance level expected under the null hypothesis that attributes and values had no influence on the neuron’s activity. Finally, we defined the value coding index with the equation:

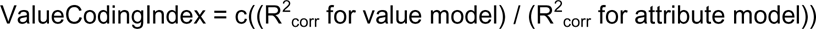

…where the function c(x) = max(0,min(1,x)) clamps the index between 0 and 1 (for rare cases where the index was slightly < 0 or > 1 such as due to small variations in the shuffling used for the shuffle correction). For some analyses, we used a signed value coding index, which was simply the value coding index multiplied by the sign of the fitted ‘Value’ weight in the value model. Importantly, because the value coding index is expressed as a ratio relative to the corrected variance explained by the attribute model, it is only meaningful and reliably estimated for neurons for which a meaningful fraction of the variance was explained by the attribute model. Therefore, we only computed this index for neurons that were strongly attribute-responsive, which we defined as neurons for which the R^2^_corr_ for the attribute model was greater than or equal to 0.03 and were fit with a significant effect for at least one attribute using the attribute model.

#### Intrinsic electrophysiological properties

Spike width was calculated as the duration between the peak and the trough of the waveform. Baseline measures of firing rate and irregularity were computed based on activity in the last 1000 ms of the ITI. We used an irregularity metric (IR) that was previously used for DRN neurons^33,34,242^, described here in brief. First, interspike intervals (ISIs) were computed. If spike(i − 1), spike(i), and spike(i + 1) occurred in this order, the duration between spike(i − 1) and spike(i) corresponds to ISI_i_; the duration between spike(i) and spike(i + 1) corresponds to ISI_i_ + 1. Second, the difference between adjacent ISIs was computed as |log(ISI_i_/ISI_i_ + 1)|.

Finally, we computed IR as the median of all such differences during the baseline window. Thus, small IR values indicate regular firing and large IR values indicate irregular firing. This measure has an advantage over traditional measures of irregularity, such as the coefficient of variation of the interspike intervals, which require a constant firing rate during the measurement period. However, for comparison, we also computed the coefficient of variation (CV) of ISIs.

## Acknowledgements

We thank Dr. Suzanne Haber and colleagues for performing serotonin transporter (SERT) staining on macaque tissue and providing us with the data. We thank Ms. Kim Kocher for excellent animal care and handling. We thank Dr. Larry Snyder for assistance with imaging. This work was supported by the National Institute of Mental Health under Award Numbers F30MH130103 to Y-YF and R01MH128344, R01MH110594, R01MH116937, and Conte Center on the Neurocircuitry of OCD MH10643 to IEM, by the Army Research Office 78259-NS-MUR to IEM, and by the McKnight Foundation award to IEM.

---

**Supplementary Table 1.**
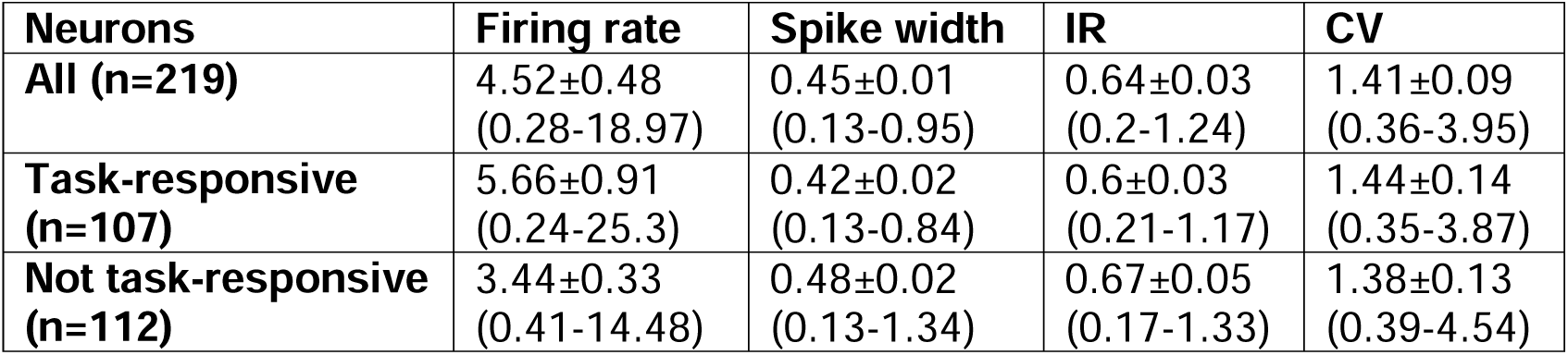
Electrophysiological characteristics of recorded DRN neurons based on task- responsiveness. Intrinsic electrophysiological properties of subsets of recorded neurons grouped by task- responsiveness. Properties were computed based on baseline activity (see Methods). Each property is shown as Mean ±1SE (95% range) for each subset of neurons. Properties did not significantly differ between groups. IR = irregularity index (see Methods), CV = coefficient of variation.

**Supplementary Table 2.** Distribution of recorded neuron locations. Mean and standard deviation of all recorded neurons’ (n=219) antero-posterior locations (AP), relative to interaural (where positive indicates anterior), and latero-medial locations (LM), relative to midline (where positive indicates to the right).

|  | AP | LM |
| --- | --- | --- |
| Mean | 3.43 | 0.06 |
| SD | 0.58 | 0.60 |

**Supplementary Figure 1.**
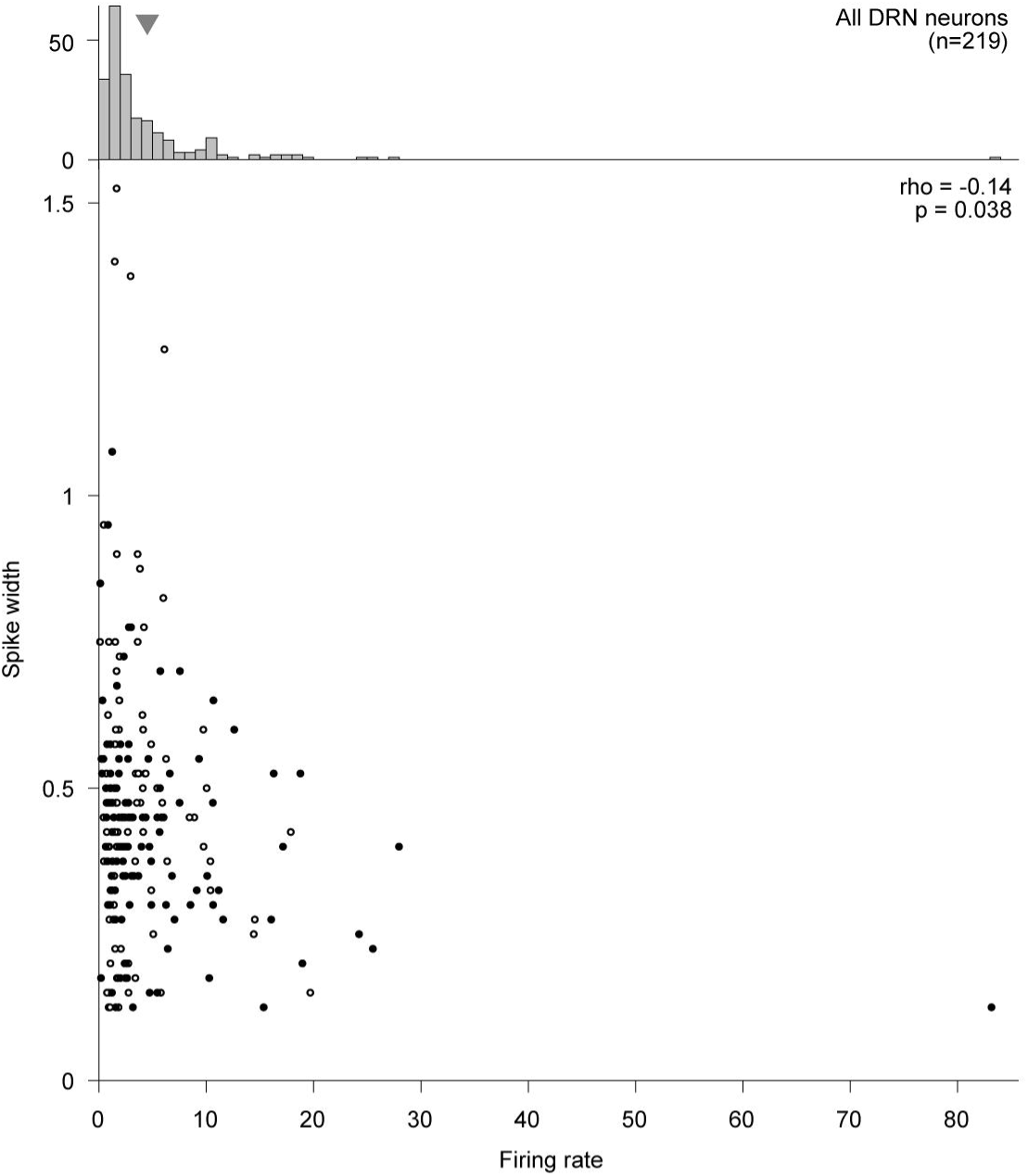
Electrophysiological characteristics of recorded DRN neurons based on task responses. Scatter plot showing spike width (ms; see Methods) vs mean baseline firing rate (Hz) of recorded neurons. Spike width was significantly negatively correlated with mean baseline firing rate (Spearman correlation, two-tailed permutation test). Filled and empty circles show neurons that were task-responsive or not, respectively. Top: Histogram showing mean baseline firing rate of recorded neurons. Marker indicates mean across neurons.

**Supplementary Figure 2.**
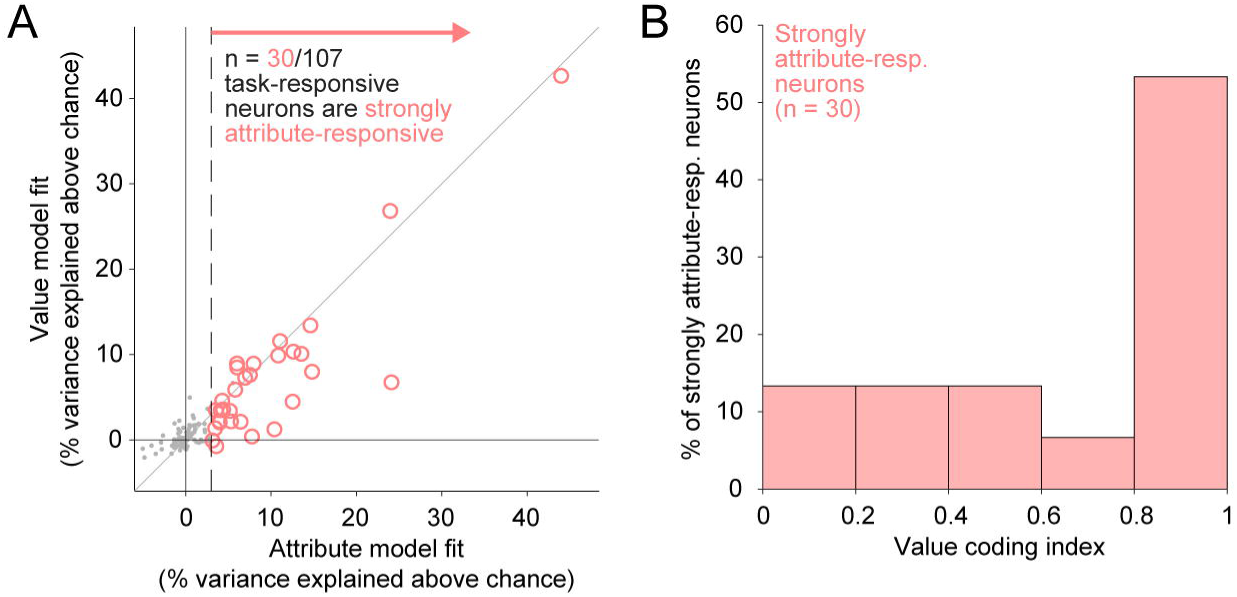
DRN neurons signal the overall subjective value of offers during multi-attribute decision-making. (A) Scatter plot showing % of variance in neuronal offer responses explained above chance (100 x R^2^_corr_) by the attribute model (x-axis) or value model (y-axis), respectively, for each task-responsive DRN neuron. Vertical dashed line indicates the threshold (attribute model R^2^_corr_ > 0.03) for classifying cells as strongly (colored circles) or not strongly (gray points) attribute-responsive (see Methods). (B) DRN neurons exhibit a strong tendency to code attributes according to their contributions to the overall subjective value of an offer, shown by histogram of value coding indices from strongly attribute-responsive DRN neurons.

**Supplementary Figure 3.**
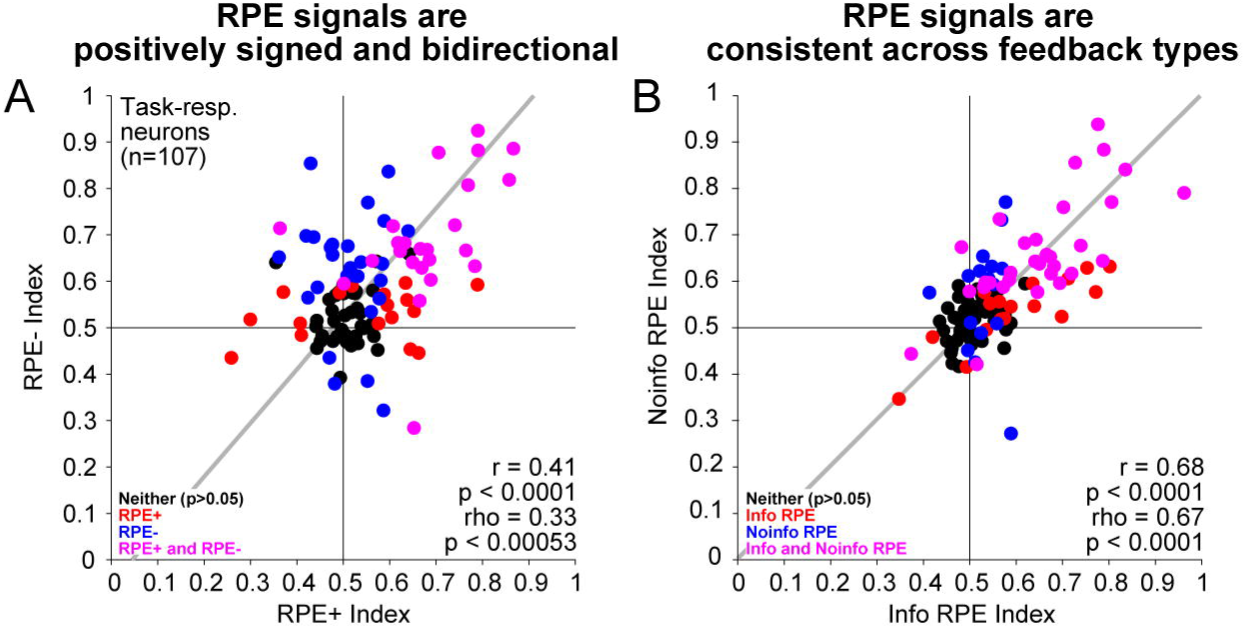
DRN neurons signal positively signed reward prediction error in response to post-decision feedback. (A) Scatter plot showing RPE+ indices vs RPE- indices (see Methods) across task-responsive neurons. Positive and negative components of RPE were strongly positively correlated (Pearson and Spearman correlations, two-tailed permutation tests), consistent with a positively signed RPE. Gray line shows Type 2 regression. Colors indicate that neither (black), the x-coordinate (red), the y-coordinate (blue), or both (magenta) are significant (p < 0.05). (B) Scatter plot showing Info RPE indices vs Noinfo RPE indices (see Methods). RPEs elicited by Info Cue and Noinfo Reveal were strongly positively correlated (Pearson and Spearman correlations, two-tailed permutation tests), consistent with a signal that generalizes between trial and feedback types. Gray line shows Type 2 regression. Colors indicate that neither (black), the x-coordinate (red), the y-coordinate (blue), or both (magenta) are significant (p < 0.05).

**Supplementary Figure 4.**
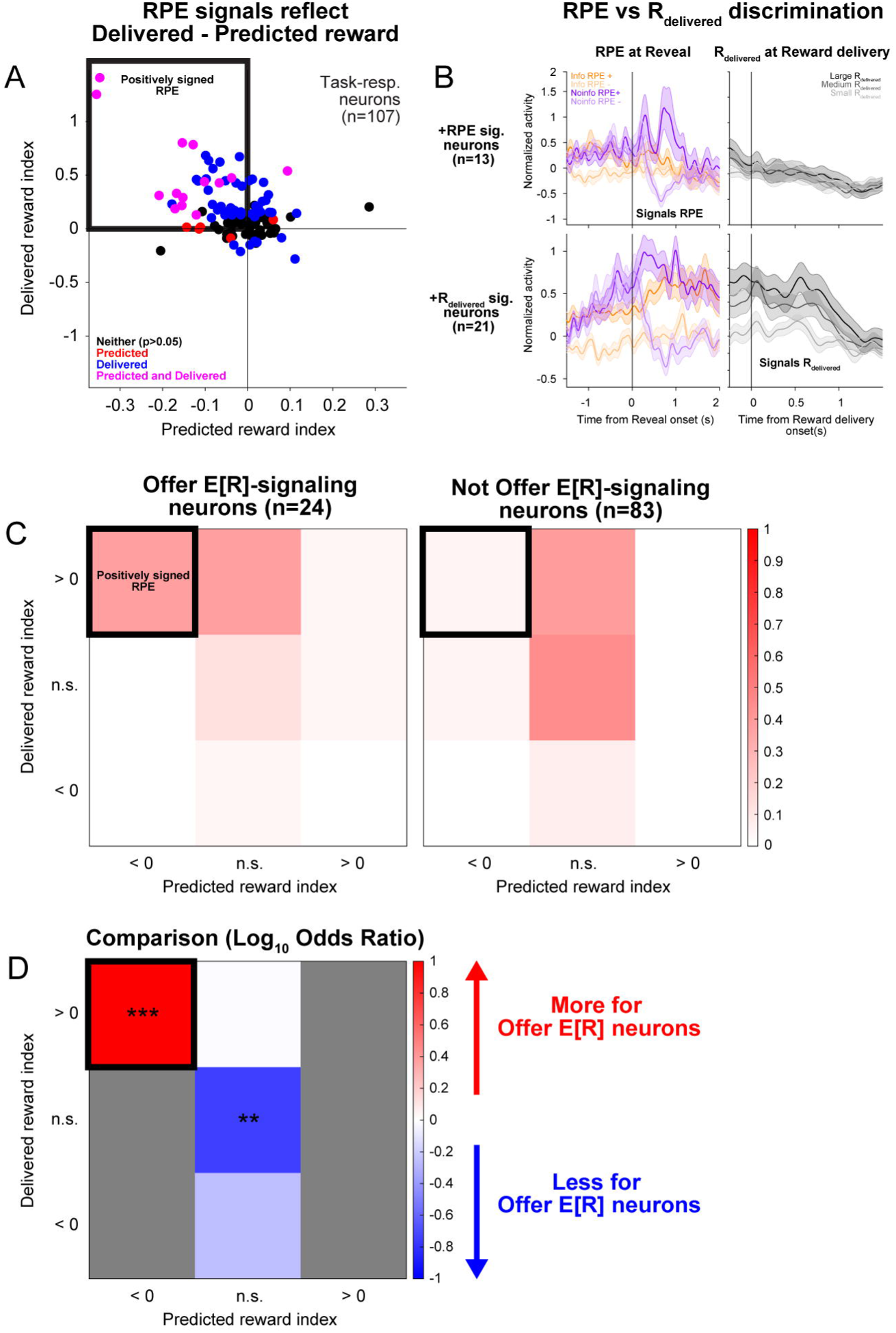
DRN reward prediction error signals reflect difference between delivered and predicted rewards. (A) Scatter plot showing delivered reward indices vs predicted reward indices (see Methods). Neurons were concentrated in the upper-left quadrant, consistent with signals that positively reflect delivered reward amount and negatively reflect predicted reward amount (i.e. a positively signed RPE). Colors indicate that neither (black), the x-coordinate (red), the y-coordinate (blue), or both (magenta) are significant (p < 0.05). (B) PSTHs showing average normalized activity in response to Reveals on Info or Noinfo trials with positive or negative RPEs (‘RPE at Reveal’, left) or that delivered Small, Medium, or Large amounts of reward (R_delivered_) (‘R_delivered_ at Reward delivery’, right) for neurons with significantly positive delivered and significantly negative predicted reward indices, (‘+RPE sig. neurons’, top) or those with significantly positive delivered reward indices but non-significant predicted reward indices that were at least tan(10°)-fold weaker in magnitude than the corresponding delivered reward indices (i.e. within 10° of the positive y-axis of the scatter plot shown in A) (‘+R_delivered_ sig. neurons’, bottom). +RPE sig. neurons were excited by positive RPEs and inhibited by negative RPEs, in response to the Reveal event on Noinfo trials, and did not phasically modulate firing in response to the Reveal event on Info trials, when feedback had already been delivered by the Cue; but these cells did not discriminate R_delivered_ following reward delivery. Conversely, while +R_delivered_ neurons were more excited by positive vs negative RPEs in response to the Reveal event on Noinfo trials, they responded similarly on Info trials, for which Reveals should not elicit RPEs; also, these neurons discriminated R_delivered_ following reward delivery. (C) Heatmaps showing proportions of Offer E[R]-signaling (left) or not Offer E[R]- signaling (right) neurons in each of 3×3 categories: (significantly positive, not significant, or significantly negative delivered reward indices)x(significantly positive, not significant, or significantly negative predicted reward indices). Neurons that signal positively signed RPEs should appear in the top left (highlighted in black). (D) Heatmap showing log_10_ odds ratio of Offer E[R]-signaling vs not Offer E[R]-signaling neurons for each of the 3×3 categories shown in (C). Offer E[R]-signaling neurons were significantly more likely to have significantly positive delivered reward indices and significantly negative predicted reward indices, consistent with a positively signed RPE signal, and significantly less likely to have neither index be significant, while none of the other proportions significantly differed (Two-tailed Fisher’s exact test; ** p<0.01, *** p < 0.001). Gray squares represent undefined values due to no neurons being in those categories in one or both groups of neurons.

**Supplementary Figure 5.**
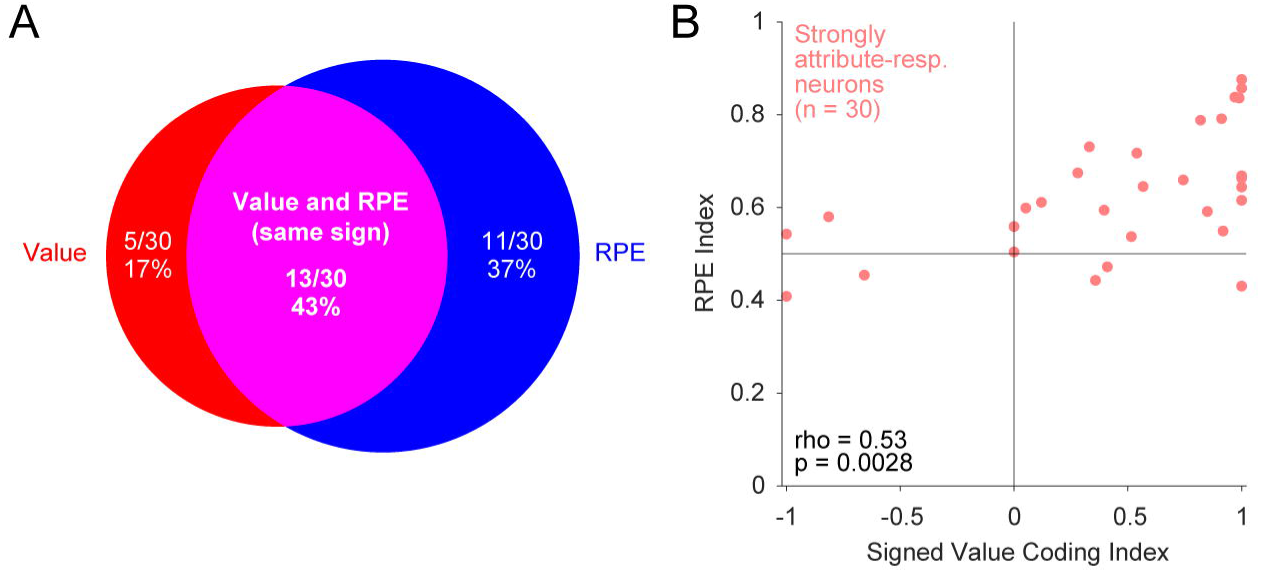
DRN neurons that signal offer value during decision-making commonly signal reward prediction errors after decision-making. (A) Modified Venn diagram showing overlap between strongly attribute-responsive neurons with high value coding indices (>0.6) (left, red) and significant RPE indices (right, blue), where the intersection includes only neurons whose value coding indices were high, RPE indices were significant, and signed value coding indices and RPE indices had the same sign (>0 and >0.5, respectively). (B) Scatter plot showing RPE indices vs signed value coding indices. RPE indices and signed value coding indices were significantly positively correlated (Spearman correlation, two-tailed permutation test).

